# An experimental demonstration of ensemble epistasis in the lac repressor

**DOI:** 10.1101/2022.10.14.512271

**Authors:** Anneliese J. Morrison, Michael J. Harms

**Affiliations:** Department of Chemistry and Biochemistry, University of Oregon, Eugene, OR 97403; Institute for Molecular Biology, University of Oregon, Eugene, OR 97403

## Abstract

Epistatic, non-additive, interactions between mutations reveal the functional architecture of living systems, strongly shape evolution, and present a difficult challenge for bioengineers. Interpreting and modeling epistasis requires knowledge of the mechanisms that bring it about. We recently argued that “ensemble epistasis” could be a generic mechanism for epistasis between mutations introduced into a single macromolecule. Because proteins exist as ensembles of interconverting conformations, a mutation could induce epistasis by subtly altering ensemble composition and thus the effects of subsequent mutations. Here we show experimentally that the thermodynamic ensemble does indeed yield high magnitude epistasis in the lac repressor. We observed two- and three-way epistasis in DNA binding, with magnitudes as large or larger than the individual effects of mutations. This biophysical effect propagated to substantial epistasis in gene expression *in vivo*. As predicted in previous theoretical work, IPTG concentration tunes the magnitude of ensemble epistasis. Further, our observations could all be captured with a rigorous mathematical model of the lac repressor ensemble. Given that conformational ensembles are unavoidable features of macromolecules, we expect this is a ubiquitous and underappreciated cause of intramolecular epistasis.

## INTRODUCTION

Non-additivity between mutations—epistasis—is ubiquitous throughout biology. It occurs across all scales of biological complexity, from individual macromolecules to entire microbial communities ^1–11^. Intramolecular epistasis between mutations introduced into a single macromolecule is particularly important, strongly shaping molecular evolution and complicating biological engineering efforts^2,12–17^. Further, intramolecular epistasis can provide important insights into macromolecular sequence-function relationships^1,3,6–9,13^. Understanding the origins of intramolecular epistasis is thus important for dissecting molecular function, improving bioengineering outcomes, and understanding how macromolecules evolve.

We recently argued that intramolecular epistasis can, theoretically at least, arise from the conformational ensembles of macromolecules ^15,18^. The conformational ensemble is the set of interchanging structural conformations a macromolecule adopts under a specific environmental condition ^19,20^. For example, the ensemble of the lac repressor—a bacterial transcription factor— consists of a conformation with high DNA affinity, a conformation with low DNA affinity, and those conformations with DNA and/or effector molecules bound (Fig 1A). Changes in effector concentration alter the relative population of each conformation within the ensemble, thus tuning transcriptional output. Conformational ensembles are generic features of biomolecules that play key roles in signaling^21^, catalysis and enzyme promiscuity ^22–25^, molecular recognition ^26,27^, and environmental sensing^28^.

**Figure 1.**
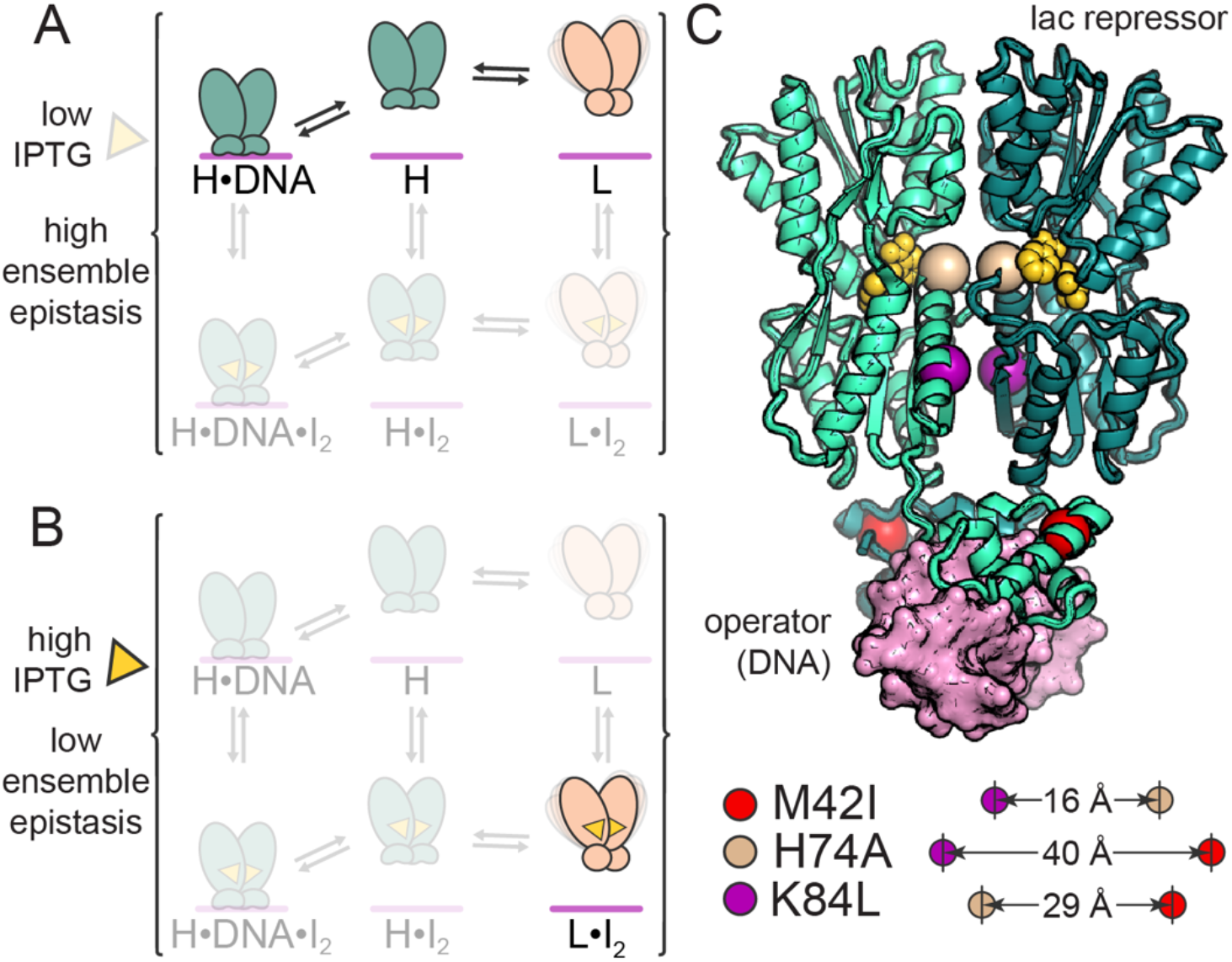
Effector concentration tunes the lac repressor ensemble. The lac repressor has two conformations: *H* (teal) has high affinity for operator (pink bar) and low affinity for IPTG (yellow triangle), *L* (peach) has low affinity for operator and high affinity for IPTG. A) At low IPTG concentrations, we see three species: H bound to operator (H·DNA), free H, and free L. B) At high IPTG concentrations, we see only L bound to two IPTG molecules (L·I_2_). Ensemble epistasis could occur at low, but not high, IPTG concentration. C) Structure of the lac repressor dimer (green cartoons) bound to operator DNA (pink surface) and an anti-inducer (yellow spheres). The anti-inducer binds at the IPTG binding site. The structure is in the H conformation (PDB 1EFA^29^). The C_α_ carbons for the sites mutated in this study are shown as spheres: M42 (red), H74 (wheat), and K84 (purple). The icons below show the distances between these C_α_ carbons in the structure.

Using our theoretical model, we predicted that ensembles would lead to epistasis when two conditions were met: 1) The macromolecule populates more than one conformation and 2) Mutations have different effects on at least two of the conformations ^18^. This leads to epistasis because the first mutation re-distributes the population of conformations in the ensemble, changing the energetic effect of a second mutation. If, for example, the first mutation stabilized a key conformation in the ensemble, it could amplify the effect of the second mutation that alters the functional output of that conformation.

The prevalence of ensemble epistasis in real macromolecules remains unknown. Our theoretical work raised three important questions: Can we detect ensemble epistasis experimentally? What is its magnitude in real macromolecules? Does epistasis at this microscopic, energetic scale propagate to meaningful epistasis *in vivo*?

To tackle these questions, we took advantage of a prediction from our theoretical work that allosteric macromolecules would be particularly prone to ensemble epistasis. This is because they populate multiple conformations as part of their function. Further, we predicted that ensemble epistasis would depend on allosteric effector concentration. At some effector concentrations, we would expect multiple conformations and thus ensemble epistasis (Fig 1A); at other effector concentrations, we would expect a single dominant conformation and thus little ensemble epistasis (Fig 1B). These considerations suggest we might experimentally detect ensemble epistasis by looking for effector-dependent epistasis in an allosteric macromolecule.

Here, we measured the magnitude of ensemble epistasis for combinations of mutations in the lac repressor both *in vivo* and *in vitro*. We observed effector-dependent ensemble epistasis for all four mutant cycles. We were able to reproduce the *in vivo* epistasis with a biophysical model of the lac repressor ensemble. This reveals that epistasis at the energetic level of the ensemble can indeed alter biological outputs. Our findings illustrate that epistasis arising from the conformational ensemble is a detectable—and likely prevalent—mechanism for epistasis in proteins.

## RESULTS

### The lac repressor has a tunable conformational ensemble

We selected the lac repressor as a model system to look for evidence of ensemble epistasis (Fig 1A,B). The lac repressor has a well-characterized conformational ensemble^30^, a measurable biological transcriptional output, and was previously shown to exhibit effector-dependent epistasis^29,31–42^. The repressor exists in two main conformations, H and L (for <u>H</u>igh and <u>L</u>ow DNA affinity). Relative to one another, H has high affinity for DNA but low affinity for the effector IPTG; L has high affinity for IPTG but low affinity for DNA. IPTG binding shifts the ensemble to favor the L conformation, causing the repressor to dissociate from DNA. We can experimentally manipulate the relative populations of the conformations by changing the concentration of IPTG. Based on previous work^30^, multiple conformations (H·DNA, H, and L) are populated with no IPTG present (Fig 1A) while a single conformation (L·I_2_) is populated at high IPTG concentrations (Fig 1B). Because ensemble epistasis requires multiple conformations, we expect ensemble epistasis is possible at low, but not high, IPTG concentrations.

We selected three well-characterized mutations to investigate ensemble epistasis in this protein: M42I, H74A, and K84L ^43–46^. M42I stabilizes operator bound conformations, H74A stabilizes effector bound conformations, and K84L stabilizes effector and operator bound conformations ^43,45–48^. These mutations are between 16 and 40 Å apart in the structure (Fig 1C). To our knowledge, these mutations have no known epistatic interactions between them. To potentially reveal both pairwise and three-way interactions between mutations, we studied eight lac repressor constructs: wildtype, three single mutants (m42I, h74A, k84L), three double mutants (m42I/h74A, m42I/k84, h74A/k84L), and the triple mutant (m42I/h74A/k84L). For clarity, we will use lowercase and uppercase letters to denote the wildtype and mutant states respectively. We will also refer to each variant by its three-letter genotype. Thus, the wildtype is “mhk,” the triple mutant is “IAL,” and the variant with the H74A mutation is “mAk.”

### *We observe IPTG-dependent epistasis* in vivo

We first tested for IPTG-dependent epistasis *in vivo* using a gene reporter assay. We placed YFP under control of the lac operon and then measured the ability of different lac repressor variants to control YFP expression in an IPTG-dependent manner ^30^. Fig 2A-D shows the induction curves for all eight variants, organized by mutant cycle. YFP expression, expressed relative to constitutively expressed YFP control, ranged from 0.0 to 0.6. For the wildtype mhk construct, the relative YFP rose from 0.0 to 0.5 between 1 μM and 1 mM IPTG. Six of the eight variants gave detectable IPTG-dependent changes in YFP expression. Three genotypes (mAk, mhL, and mAL) were “leaky”, allowing YFP transcription at low IPTG concentrations. All genotypes containing the k84L mutation had severely diminished induction responses.

**Figure 2.**
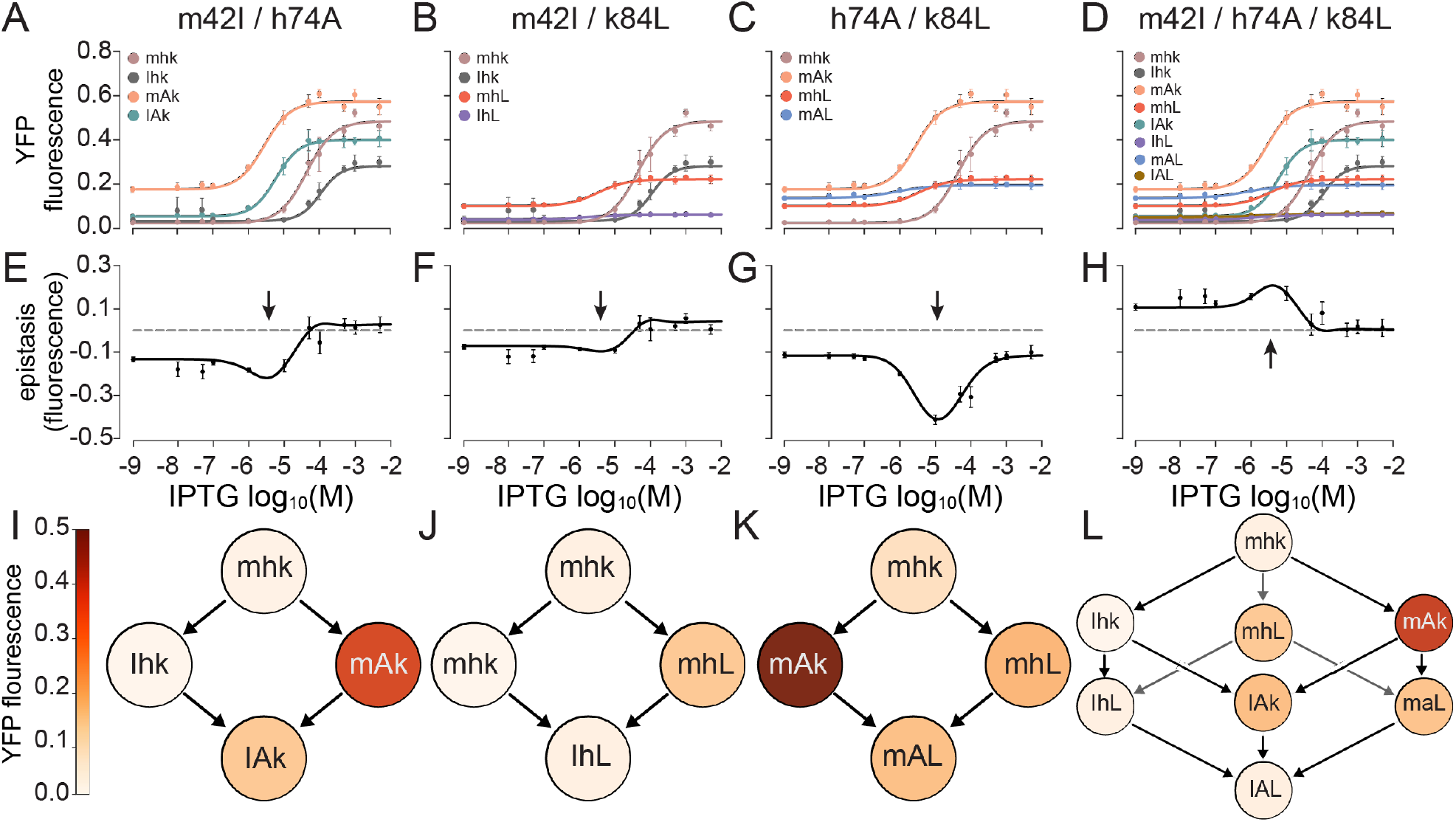
Mutations exhibit IPTG-dependent epistasis *in vivo*. A-D) Induction curves for lac repressor genotypes. Plots show normalized YFP expression versus IPTG concentration. Each series represents a single genotype, labeled on the plot. Each point is the mean of three or more biological replicates, with their standard deviation shown as error bars. Lines represent a Hill model fit to the averaged data. E-H) Epistasis as a function of IPTG for the induction curves above. Circles represent the magnitude of epistasis averaged over 100 bootstrapped datasets extracted from the uncertainty in the experimental data. Error bars are the standard deviation. Lines show epistasis in the Hill model values (i.e., epistasis calculated using the smooth lines in panels A-D). For the three-way mutant cycle, the epistasis shown is third-order epistasis. I-L) YFP expression for each genotype in the four mutant cycles at the IPTG concentration that gives maximum epistasis. This concentration is indicated with arrows in E-H. The color bar indicates normalized YFP expression.

We next calculated epistasis as a function of IPTG concentration for these mutant cycles. For each mutant pair (a→A/b→B), we calculated epistasis as the difference in the effect of mutation a→A in the wildtype (ab) versus mutant (aB) genetic backgrounds. For the m42I and h74A mutants, this is given by:

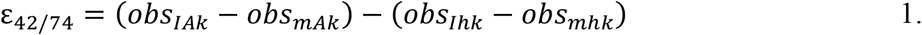

where *obs* is the YFP expression at a given IPTG concentration and the subscripts indicate the genotype. Analogous expressions were used for the other mutant pairs. For the three-way mutant, we defined high-order epistasis as the difference between the observed YFP expression of the triple mutant and the predicted expression given the individual and pairwise effects of the three mutations^17^. This is given by:

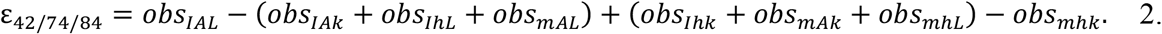

Fig 2E-H show epistasis as a function of IPTG concentration for the four cycles. At low IPTG concentrations (< 10^−7^ M), we see a moderate amount of epistasis for all four mutant cycles. At intermediate IPTG concentrations (10^−7^ M < [IPTG] < 10^−3^), we see a peak in the epistatic magnitude. For the pairwise cycles, the epistasis is negative, meaning the effect of the combined mutations on induction is less than expected given their additive effects. For the three-way cycle, the epistasis is positive, meaning the effect of the combined mutations on induction is higher than expected given their additive and pairwise effects. For all cycles, the peak coincides with the IPTG concentrations over which the switch-like induction response occurs (Fig 2A-D). At high IPTG concentrations (> 10^−3^ M), we see a decrease in the magnitude of epistasis, which approaches zero in three out of the four mutant cycles. The maximum epistasis for each set of genotypes is shown as a mutant cycle in Fig 2I-L.

The magnitude of the epistasis in these curves is substantial. The total induction response of these transcription factors goes from 0.0 to 0.6 (Fig 2A-D). We observed epistasis as high as - 0.4 (e.g., h74A/k84L cycle at 10^−5^ M IPTG; Fig 2G): the epistasis is ∼70% as large as the total dynamic range for the wildtype repressor. These large-magnitude effects are not limited to the pairwise epistatic curves: the high-order epistasis from the three-way mutant has a maximum magnitude near +0.2 (Fig 2H, right)—35% of the magnitude of the total induction of the wildtype repressor.

### A thermodynamic linkage model captures the conformational ensemble

The IPTG-dependent epistasis we observed *in vivo* is consistent with ensemble epistasis— peaking where we might expect many populated conformations and dropping to zero where we expect only one. We next wanted to see if this was indeed the mechanism leading to IPTG-dependent epistasis. To do so, we characterized the repressor ensemble *in vitro* (Fig 3) and then attempted to reproduce our *in vivo* measurements with this mechanistic information. We described the ensemble using a previously published mathematical model of the lac repressor consisting of twelve species whose populations are determined by three concentrations (total repressor, total IPTG, and total DNA operator) and five equilibrium constants (K_HL_, K_HI_, K_LI_, K_H-DNA_, K_L-DNA_) (Figure 3A) ^30^.

**Figure 3:**
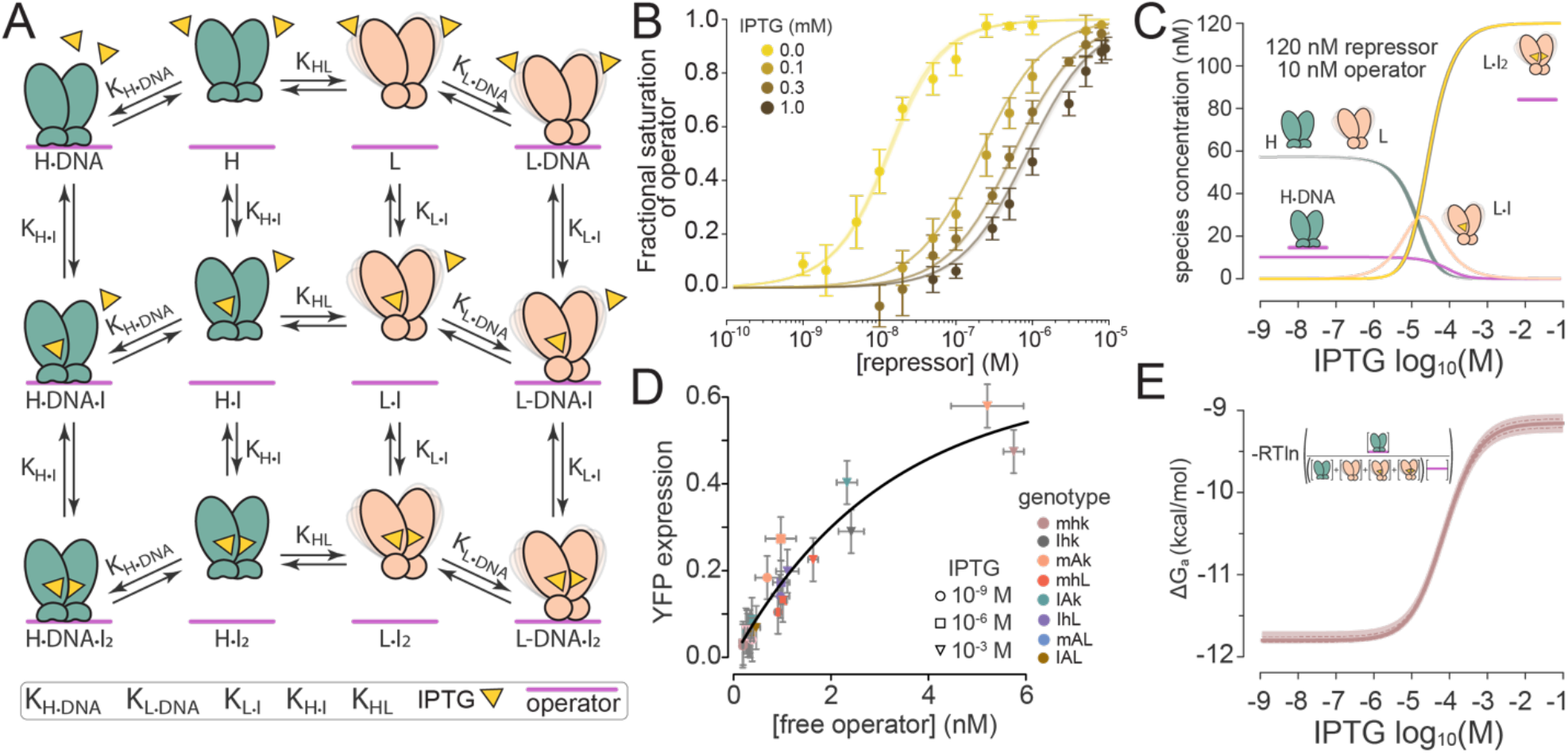
The molecular ensemble of the lac repressor can be described with a mathematical model. A) Model of the lac repressor ensemble with twelve species and five equilibrium constants ^30^. The icons and equilibrium constants are labeled on the figure. B) Binding of the wildtype mhk repressor to operator DNA *in vitro*, as measured as measured by fluorescence anisotropy. Plot shows fractional saturation of operator versus repressor concentration. Points and error bars indicate the average and standard deviation of at least three biological replicates. Clouds of lines show sampled model parameter sets extracted from a global Bayesian MCMC analysis. The color of each dataset corresponds to the IPTG concentration as indicated on the figure. Note: Each IPTG series included a 0 M repressor sample that is not shown on the plot. C) Concentrations of molecular species from panel A calculated using the model parameters extracted from panel B. Only species with concentrations above 1 nM are plotted. Shaded area around each curve indicates the 95% credibility interval taken from the MCMC samples. The “H” and “L” curves exactly overlap. Calculations were done using 120 nM total repressor, 10 nM total operator, and variable IPTG concentrations. D) *In vivo* YFP expression versus free operator concentration calculated using the MCMC model parameters. Point colors indicate genotype; point symbols indicate IPTG concentration. Line indicates a least-squares fit of the model A×K×[IPTG]/(1 + K×[IPTG]) to the data. Errors are standard deviations of biological replicates (YFP expression) and 95% credibility interval (operator concentration). E) ΔG_a_ versus IPTG concentration calculated from the concentrations of the species in panel C. Heavy line is the median of the MCMC samples; dashed lines are the standard deviation; the shaded region indicates the 95% credibility interval.

We purified each variant and used fluorescence anisotropy to measure its ability to bind operator DNA at ∼6-10 repressor concentrations and four IPTG concentrations (Fig 3B, Fig S1-S8). We used a Bayesian Markov-Chain Monte Carlo (MCMC) strategy to sample model parameters consistent with all observations for each protein. In preliminary analyses, we found that not all model parameters were well constrained by the experimental data and that some covaried strongly with one another (Fig S9). This matches what was observed by the group that developed the model ^30^. To improve the fits, we made three modifications to the model. First, we set K_L-DNA_ to 1 × 10^−10^ M, following previous observations that the repressor has weak affinity for DNA in the L conformation ^30^. Second, because our estimates for K_HL_ and K_H·DNA_ always co-varied (Fig S9-S13), we reduced this to a single parameter K_DNA_ (K_H·DNA_/K_HL_). This parameter is the effective protein/DNA binding constant that incorporates the energetic penalty required to shift the protein from the L to H conformation. Third, we simultaneously fit the model against our *in vitro* binding data and our *in vivo* induction curves. This required the addition of a nuisance parameter accounting for the small difference in operator affinity *in vitro* and *in vivo*. An Akaike Information Criterion (AIC) test strongly supported our final model relative to models with fewer or greater numbers of parameters (p = 10^−64^; see methods for details).

Fig 3B shows the binding measurements and binding curves calculated from the MCMC samples for the wildtype mhk variant. The model captures the experimental observations well, with low, random residuals across all 40 experimental conditions (Fig 3B, S1). Further, the model parameters are well constrained by the data. (Here, and throughout the rest of the text, we report these values as standard state free energies, where ΔG° = -RTln(K)). For the mhk variant, the final model parameters were ΔG°_DNA_ = 0.92 [0.70,1.12], ΔG°_H·I_ = 7.76 [7.84,7.76], and ΔG°_L·I_ = 5.98 [5.85,6.11] kcal/mol. We report the median value and the lower and upper bounds of the 95% credibility interval in brackets. Our parameter estimates the values reported by the group that developed the model (ΔG°_DNA_ = 1.6 ± 0.5, ΔG°_H·I_ = 8.0 ± 0.3, and ΔG°_L·I_ = 6.5 ± 0.3 kcal/mol). All fit parameters are reported in Table S1.

We next used the extracted model parameters to calculate the concentration of the species in the linked equilibria (Fig 3C). For these calculations, we used total repressor and operator concentrations of 120 nM and 10 nM, respectively, matching our estimated concentrations of repressor and operator *in vivo* (see methods). We varied the total IPTG concentration between 1 nM and 10 mM IPTG. Over these concentration ranges, only five species were appreciably populated by the mhk variant. At 1 nM IPTG, the ensemble consists of H, L, and H·DNA. At ∼0.1 mM IPTG, the ensemble consists of H, L, H·DNA, L·I, and L·I_2_. Finally, at 10 mM IPTG, the ensemble consists solely of L·I_2_ (Fig 3C).

To validate the consistency between our model and our *in vivo* measurements across all genotypes, we used the model to calculate the concentration of free operator as a function of IPTG for all eight lac repressor variants. We then used this concentration to predict relative YFP expression *in vivo*. Figure 3D shows YFP expression versus operator concentration estimated from the thermodynamic model for all variants and three IPTG concentrations. We see a strong correlation between the two values. As expected, this function saturates with increasing free operator concentrations. At the highest operator concentrations, other cellular factors besides the number of operators exposed controls YFP yield. A simple saturation model explains this relationship with an R^2^ of 95.7.

### *IPTG-epistasis occurs* in vitro

We next asked whether the IPTG-dependent epistasis we observed *in vivo* arose from the lac repressor itself or by interaction with other factor(s) in the cell. To do so, we attempted to reproduce the *in vivo* epistasis using our mathematical model of the conformational ensemble. We used the apparent binding free energy of the repressor/DNA complex (ΔG_a_) as our observable. We selected this observable because free energies are on a linear scale and are thus appropriate for epistatic analyses^49^. ΔG_a_ can be calculated from the ensemble by:

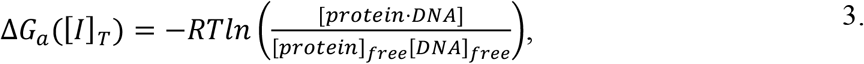

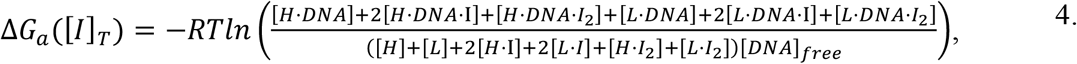

where R is the gas constant, T is the temperature, [I]_T_ is the total IPTG concentration, and the remaining concentrations are the equilibrium concentrations of the molecular species shown in Fig 3A. Fig 3E shows how ΔG_a_ changes for the mhk variant as a function of IPTG. There is a 2.5 kcal/mol change in affinity for DNA between 1 nM and 10 mM IPTG, reflecting the observed shift from the H·DNA to the L·I_2_ species. ΔG_a_ strongly correlates with the magnitude of YFP induction across all eight variants (Fig S14).

Figure 4A-D shows ΔG_a_ vs. [IPTG] for the four mutant cycles under investigation. Between 1 nM and 10 mM IPTG, we observed changes in ΔG_a_ as large as +2.5 kcal/mol (variant mhk) and as small as 0 kcal/mol (variant IAL). These curves closely match those observed for the *in vivo* induction curves (Figure 2A-D). We next calculated epistasis in ΔG_a_ as a function of IPTG using equations 1 and 2. For all four cycles, we observed moderate epistasis at low IPTG concentration, a peak at the intermediate IPTG concentration, and no epistasis at high IPTG concentration (black curves, Fig 4E-H). These curves have highly similar shapes to those extracted from YFP expression (blue curves, Fig 4E-H). This verifies that the IPTG-dependent epistasis in induction we observed is encoded by the protein itself, not interactions with other factors in the cell.

**Figure 4.**
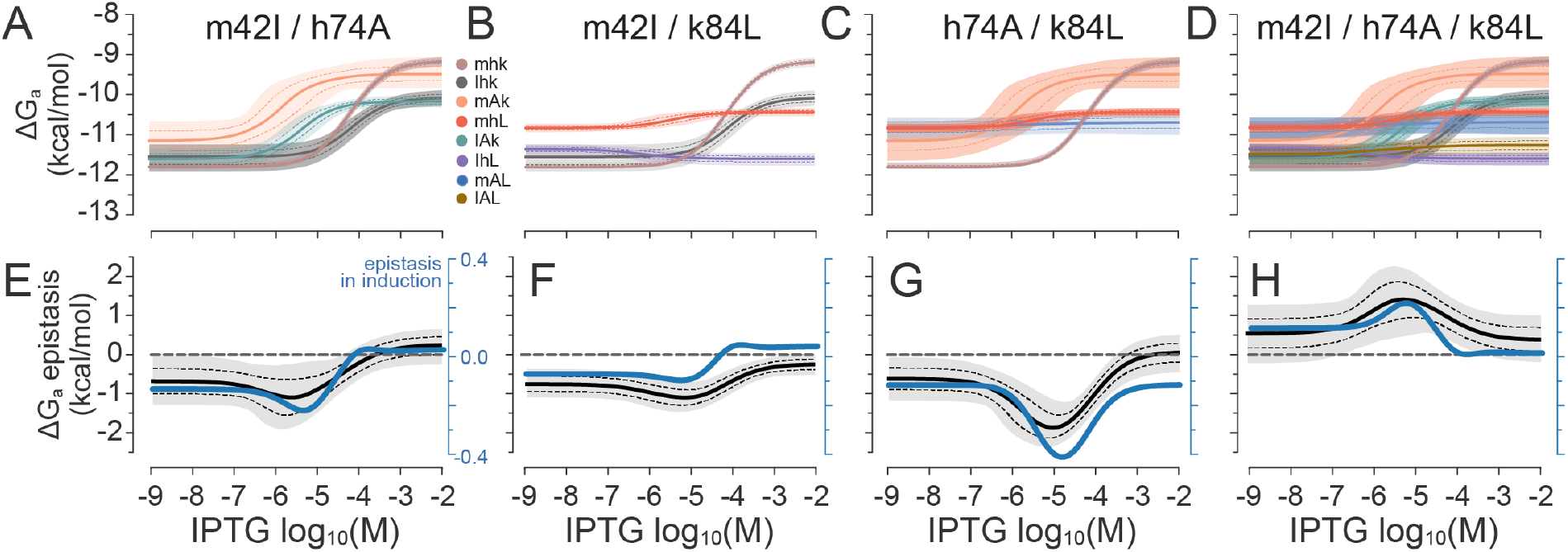
*In vivo* IPTG-dependent epistasis can be reproduced by the ensemble model. A-D) Apparent DNA binding affinity (ΔG_a_) versus IPTG concentration for the four mutant cycles. Colors indicate genotype as shown on the plot, matching those from Figure 2. The heavy lines are the medians of MCMC samples; the dashed lines are the standard deviations; the shaded regions indicate the 95% credibility interval. E-H) Epistasis in ΔG_a_ calculated for the mutant cycles in A-D (black lines and gray shaded region). The epistasis in induction from Figure 2E-H is reproduced in blue to facilitate comparison.

### Epistasis arises from redistributing the conformational ensemble

Ensemble epistasis occurs when a mutation redistributes the conformational ensemble, thereby changing the effect a subsequent mutation. While our results are consistent with ensemble epistasis, one other possible source of epistasis in the system would be non-additive effects of mutations on ΔG°_DNA_, ΔG°_H·I_, and/or ΔG° _L·I_. To test for this possibility, we determined the epistatic interactions between mutations in DNA affinity (ΔG°_DNA_; Fig 5A-C) and in the difference in IPTG affinity between the H and L conformations (ΔG°_H·I_ - ΔG° _L·I_; Fig 5D-F).

**Figure 5.**
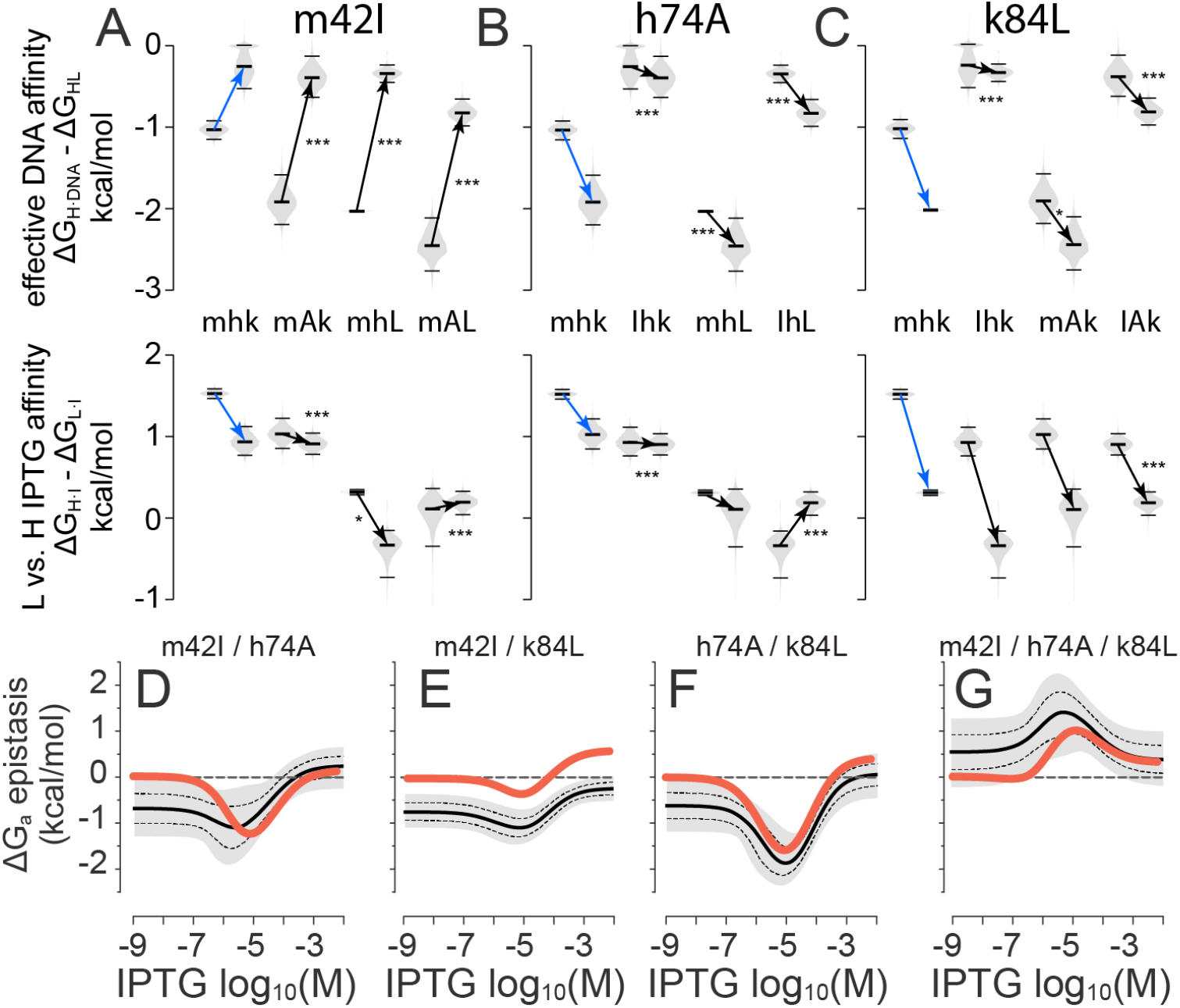
IPTG-dependent epistasis is not due to epistasis in equilibrium constants. A-E) Effects of m42I, h74A, and k84L on effective DNA affinity (ΔG°_DNA_) and the difference in the IPTG affinity between the H and L conformations (ΔG°_H·I_ – ΔG° _L·I_). Violin plots show energies extracted from the MCMC samples; horizontal lines indicate the median and 95% credibility interval. Paired violin plots show the effect of the mutation in the genetic backgrounds indicated in the center of each plot. Arrows connect the medians for the background and mutant genotypes. A difference in arrow slope between the same mutant in different genetic backgrounds indicates epistasis. The effects of the mutations in the mhk background are shown as blue arrows. Stars indicate mutations whose effect differs in the specified background versus the mhk background: p < 0.01 (*), p < 0.001 (**), and p < 0.0001 (***). D-G) Epistasis in ΔG_a_ for the mutant cycles calculated using additive effects of each mutation on ΔG°_DNA_, ΔG°_H·I_, and ΔG°_L·I_ (red lines). For comparison, epistasis using the MCMC sampled fit parameters is reproduced from Figure 4E-H (gray region; black lines).

We plotted the effect of each mutation on these energy terms across all genetic backgrounds (Fig 5A-F). All mutations exhibited epistasis in both energy terms in at least one background. m42I, for example, decreases effective DNA affinity by 0.5 kcal/mol in the mhk background but by ∼1.6 kcal/mol in all other backgrounds (Fig 5A). m42I also decreases the difference in IPTG affinity between H and L by 0.8 kcal/mol in the mhk background but has no effect in the mAk or mAL backgrounds (Fig 5D). Similar observations hold for all other mutations. Although most of the epistatic interactions were changes in magnitude rather than sign, h74A exhibited sign epistasis: h74A lowered the difference in IPTG affinity between H and L by 0.7 kcal/mol in the mhk background but increased this difference by 0.6 kcal/mol in the IhL background (Fig 5E).

To separate the contribution of ensemble epistasis from epistasis at the level of individual energy terms, we generated ensembles in which all mutations had additive energetic effects. We modeled each energy term in each genotype by summing the effects of the included mutations in the mhk background. For example, the additive ΔG°_DNA_ term for the mAL genotype was ΔG°_DNA,mhk_ + (ΔG°_DNA,mAk_ - ΔG°_DNA,mhk_) + (ΔG°_DNA,mhL_ - ΔG°_DNA,mhk_). The additive energies for each variant are given in Table S1.

We then calculated epistasis in ΔG_a_ as a function of IPTG for these additive ensembles. All mutant cycles exhibited peaks in epistasis as a function of IPTG, even using additive ensembles (Fig 5G-I). Using additive mutations did not change the sign or magnitude of the IPTG-dependent epistasis: for all cycles, the basic shape of the curve was unchanged (Fig 5G-I). The additive curves did, however, systematically differ from the non-additive curves. At low IPTG concentrations, the original cycles curves had epistasis that was not present in the additive cycles. Epistasis at low IPTG concentration is mediated at the level of equilibrium constants, not redistribution of the ensemble. There were other intriguing differences. The additive m42I/k84L and h74A/k84L cycles were globally ∼1 kcal/mol lower than their non-additive counterparts, suggesting an IPTG-independent epistatic effect in equilibrium constants. This contrasts with the m42I/h74A and m42I/h74A/k84L cycles: the non-additive and additive cycles only differed at the lowest IPTG concentrations.

Overall, these observations reveal that epistasis in the lac repressor is a combination of epistasis within individual energy terms and the redistribution of states in the ensemble. More specifically, the IPTG-dependent peak in epistasis observed for these cycles arises from the ensemble. (This matches our theoretical predictions^18^.) Epistasis at the level of equilibrium constants, in contrast, manifests as offsets to the ensemble-induced curve.

### In vivo *epistasis can be understood using the ensemble*

Finally, we sought to understand the biophysical mechanisms through which the ensemble led to epistasis (Fig 6). We characterized our pairwise mutant cycles at three levels—energetics, the ensemble, and induction—allowing us to explain epistasis in the biological phenotype in terms of the populated species (Fig 6A). For all cycles, we analyzed epistasis at 10 μM IPTG, corresponding to the peak in epistasis observed both *in vivo* and *in vitro* (Fig 4E-H). To isolate epistasis arising from the ensemble, we treated the energetic effects of each mutation additively.

**Figure 6.**
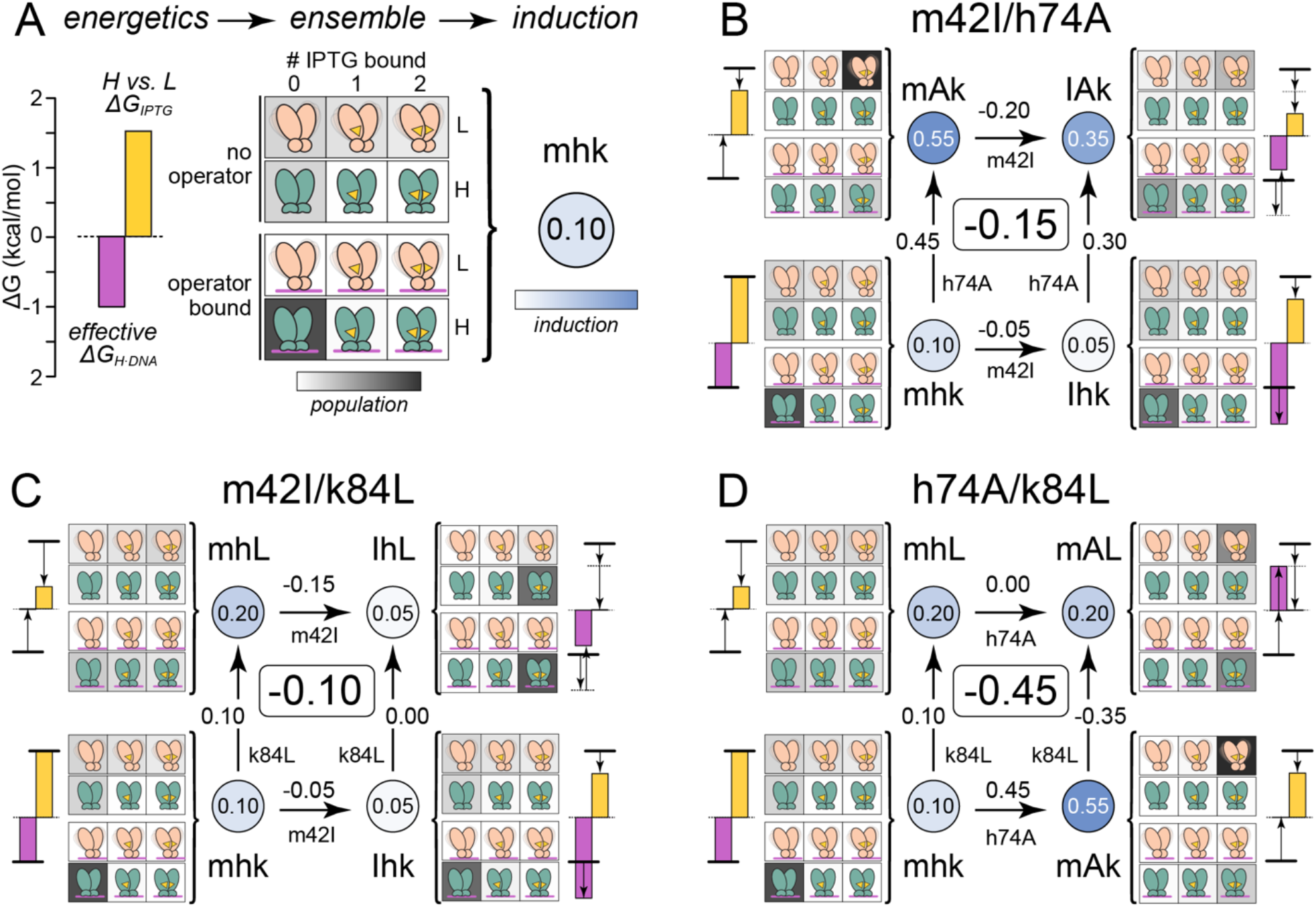
Epistasis arises from redistributing the ensemble. A) Energetics, ensemble, and induction for the mhk genotype at 10 μM IPTG. Bars show effective DNA affinity (pink; ΔG°_DNA_) and the difference in IPTG affinity of the H vs. L states (yellow; ΔG°_H·I_ – ΔG° _L·I_). Central panel shows the relative population of all species in the ensemble on a white-to-gray, low-to-high concentration gradient. For the top six, operator-free, panels the population gradient goes from 0-120 nM; for the bottom six, operator-bound, panels the gradient goes from 0-10 nM. The circle to the right indicates the induction level (0.10) on a white-blue gradient. B-D) Pairwise mutant cycles for m42I/h74A (B), m42I/k84L (C), and h74A/k84L (D). Genotype is shown at each corner on the cycle. Circles indicate induction using the white-blue gradient from panel A. Cycle arrows indicate mutations; the numbers on each arrow are the effect of that mutation on induction. The large, boxed number in the center of the cycle is epistasis in induction. Bar plots show energies, as on panel A. The arrows indicate the effects of the relevant mutations on those energetic levels. The population plots use the white-gray gradient show in panel A. All cycles are shown at 10 μM total IPTG.

In Fig 6B, we dissect the m42I/h74A mutant cycle. The m42I mutation increases the effective affinity of the repressor for DNA (pink bar), while slightly decreasing the difference in IPTG affinity between the H and L conformations (yellow bar). Overall, this leads to lower induction for Ihk than mhk (0.05 vs 0.10, respectively). The h74A mutation, in contrast, dramatically disrupts DNA binding (pink bar), while slightly decreasing the difference in IPTG affinity between the H and L conformations (yellow bar). This causes a large increase in induction (0.55 vs. 0.10) because the repressor has a weak interaction with the operator. When combined, however, we observe an induction of 0.35—not 0.50 as we would expect from adding the effects of these mutations on induction. The effects of the two mutations on DNA binding offset each other, while the effects of the two mutations on IPTG binding are in the same direction. This means the IAk repressor interacts with DNA slightly less well than mhk, but that IPTG also provides a less effective driving force to cause it to fall off. In net, this means the IAk repressor populates the operator bound, inhibited state more than the mAk genotype, explaining its lower overall induction.

For the m42I/k84L cycle, we see m42I completely overwhelming the positive effect of k84L on induction (Fig 6C). Alone, k84L increases induction from 0.1 to 0.2; in the m42I background, k84L has no effect on induction. This occurs because k84L both lowers the effective affinity of the repressor for DNA and decreases the difference in IPTG affinity between the H and L conformations. When combined with m42I—which increases affinity for DNA but also lowers the difference in IPTG affinity between the H and L conformations—the repressor has moderately high affinity for DNA and no difference in IPTG affinity between H and L. As a result, the repressor populates an IPTG- and DNA-bound state that represses expression, even in the presence of IPTG.

Finally, for the h74A/k84L cycle, we see an behavior intermediate between the previous two cycles. Together, h74A and k84L disrupt the difference in IPTG affinity between H and L, meaning IPTG is not effective at inducing expression. Further, both h74A and k84L disrupt DNA binding. This means the final repressor is both unresponsive to IPTG and has low DNA affinity. As a result, the repressor populates both a repressed IPTG-/DNA-bound conformation and an IPTG-, but not DNA-bound, conformation. This leads to a moderate amount of induction for the double mutant; higher than mhk, but lower than mAk.

## Discussion

We showed previously that protein conformational ensembles could theoretically give rise to epistasis^18^. In this work, we demonstrated the presence of ensemble epistasis in the lac repressor. In the introduction, we asked three questions, all of which can now be answered.

1. *Can we detect ensemble epistasis experimentally?* Yes. We predicted that ensemble epistasis in an allosteric protein would appear as effector-dependent epistasis ^18^. We observed this both *in vivo* (Fig 2A-D) and *in vitro* (Fig 5D-G). Ensemble epistasis is maximized at the IPTG concentration at which the lac repressor transitions from being bound to operator to bound by IPTG. Under these conditions, the ensemble populates the highest diversity of conformations and thus has the highest capacity for ensemble epistasis.
2. *What is the magnitude of ensemble epistasis?* Substantial. *In vitro*, we observed ensemble epistasis as large as 2 kcal/mol (Fig 5F). This is larger than the individual effects of the mutations (∼1 kcal/mol; Fig 5B-C). *In vivo*, we observed epistasis in YFP induction as that was almost as large as the total induction with addition of 1 mM IPTG (0.4 vs. 0.55; Fig 4C).
3. *Does epistasis at this microscopic, energetic scale propagate to meaningful epistasis* in vivo*?* Yes. Through careful biochemical characterization and mathematical modeling, we were able to demonstrate how subtle redistribution of conformational states in the ensemble was responsible for the observed epistasis between mutations in *in vivo* YFP expression (Fig 6).

### Ensemble epistasis is likely common in real mutant cycles

How common is ensemble epistasis? Several considerations lead us to believe it is extremely common. First, conformational ensembles are a general feature of macromolecules, and are often critical for biological function ^22–24,28^. This means there is ample opportunity for ensemble epistasis. Second, our previous computational work suggested that 27% of mutant pairs in an allosteric protein would exhibit ensemble epistasis with a magnitude at or above kT (0.6 kcal/mol)^18^. Third, despite having no previous knowledge that the mutations we selected for this study would exhibit epistasis, we observed ensemble epistasis for all mutant cycles. Although we only tried a few mutants, this hints that ensemble epistasis does not require careful selection of specific mutations and may therefore be common.

Finally, there is a growing body of literature documenting environmentally dependent epistasis ^50^. Sometimes this can be directly attributable to the ensemble, as in the case of the bacterial GB1 protein binding to ligand ^51^. In other cases, an ensemble-based mechanism seems plausible, though is not established. For example, a recent study of the evolutionary transition between an ancestral dihydrocoumarin hydrolase and a derived methyl-parathion hydrolase showed that epistatic interactions between historical mutations changed both magnitude and sign depending on the presence of different divalent metal ions ^52^. Environment-dependent epistatic interactions such as this have been observed in diverse macromolecular systems––from catalytic RNAs ^53,54^, drug resistance enzymes ^55–57^, yeast tRNAs ^58^, cis-regulatory elements ^59–61^, and transcription factors ^42,62^. Environment-dependent epistasis has been observed extensively in more complex biological systems, for instance in experimental evolution experiments ^63–65^.

### Ensemble epistasis is maximized where important functional transitions occur

The magnitude of ensemble epistasis is maximized under conditions where macromolecules transition between distinct, biologically important functions. Here, this corresponds to IPTG concentrations where many conformations are populated because the system is transitioning from the operator bound to effector bound conformations.

The coincidence of ensemble epistasis with function-switching transitions means that any protein that must switch states as part of its function is unavoidably evolving in a highly epistatic regime ^11,50,66–68^. This epistasis arises directly from conformational diversity, which has been linked to many important facets of evolutionary biology: phenotypic plasticity ^69,70^, increased rates of evolution ^71^, and increased robustness and evolvability ^68,72^. Further, ensemble epistasis may couple mutational effects to environmental changes, thus strongly shaping evolution. A recent study compared the trajectories of *E. coli rpoB* mutants under fluctuating antibiotic concentrations to those in constant antibiotic concentrations. In addition to pervasive environment-dependent epistasis, they found that there was more diversity in the solutions to survival in the fluctuating environments, indicating that there were more paths accessible, and that specific mutational trajectories were contingent upon intermediate environments, meaning that those trajectories are only accessible if an intermediate environment was encountered ^56^.

### Revealing other sources of epistasis

In addition to observing ensemble epistasis, we found epistasis that occurred at the level of individual equilibrium constants (Fig 5). To a first approximation, this form of epistasis was independent of IPTG concentration, reflecting a fixed interaction between residues. The magnitude of these epistatic effects (∼0.5 kcal/mol) was substantial, but slightly smaller than the maximum ensemble epistasis (∼1.0 kcal/mol) for each cycle. The biophysical mechanism(s) for this epistasis remain unclear and will require further investigation. One possibility is that this epistasis arises from direct, physical interactions between the residues within a specific conformation in the ensemble. These residues are 16 to 40 Å apart in the structure, so such an interaction would likely require a pathway or network of residues [yy please add https://pubmed.ncbi.nlm.nih.gov/14573864/]. If so, studying epistasis as a function of effector concentration may be a useful tool to isolate sites that interact through allosteric pathways rather than by tuning ensemble outputs.

### Conclusions

We found ensemble epistasis between all combinations of mutations studied. This biophysical effect led to observable changes in transcriptional output, revealing that ensemble epistasis at the biophysical level can lead to epistasis at the cellular and biological level. Because macromolecules unavoidably exist as ensembles of conformations, we expect this is a ubiquitous mechanism of intramolecular epistasis.

Important questions remain. How common is it? Is it coupled to other mechanisms of epistasis? We observed three-way ensemble epistasis: does it occur at even higher orders? How can we account for ensemble epistasis to gain a deeper, more predictive understanding of biology? Answering these questions should be a high priority as the field sets out to understand the mechanisms of epistasis and how this shapes protein biochemistry and evolution.

## Materials and methods

### *In vivo* induction assays

For our *in vivo* experiments, we cloned the wildtype lac repressor sequence into the PAM5087 vector (addgene plasmid #85123) downstream of the constitutive lacIq promoter using SLIC cloning ^73^. The PAM5087 vector contains the YFP gene downstream of the O1 lac operator sequence and a trc promoter. We generated a wildtype repressor construct c-terminally tagged with the mCherry protein. A 24 amino-acid rigid linker sequence (GSLAEAAAKEAAAKEAAAKAAAAS) was cloned in frame between the two genes to prevent protein-protein interactions ^74^. The mCherry-tagged wildtype lac repressor gene was then cloned into the PAM5087 vector. All eight genotypes were cloned upstream of the mCherry gene. We confirmed that the mCherry-tagged lac repressor was folded CD spectroscopy and by comparing the tagged and untagged wildtype induction curves (Fig S15). (This is consistent with previous observations that the mCherry-tagged lac repressor dimer has previously been confirmed to have repressor activity equivalent to the untagged repressor *in vivo* ^30,75^). Finally, we created a DEL construct in the PAM5087 background by introducing an N-terminal stop codon in the wildtype lac repressor protein.

Each construct was transformed into BLIM cells (Addgene bacterial strain #35609). BLIM cells are derived from BL26 Blue cells (Novagen) with both the lac operon and F’ episome, which contains the lac repressor gene, removed ^76^. LB media supplemented with 50 μg/mL kanamycin were inoculated from PAM5087 construct glycerol stocks and grown at 37°C, 250 RPM overnight. Overnight cultures were used to inoculate 10 mL cultures in pre-growth M9 media (47.8 mM Na_2_HPO_4_, 22 mM KH_2_PO_4_, 8.6 mM NaCl, 18.7 mM NH_4_Cl, 10 mM NaHCO_3_, 0.0025% (w/v) thiamine-HCl, 0.2% (w/v) Casamino Acids, 0.01X Basal Eagle Medium, 0.04% glycerol, 2 mM MgSO_4_, 1 mM CaCl_2_) supplemented with 50 μg/mL kanamycin. Pre-growth cultures were grown for 5-6 hours at 37°C, 250 RPM. Black-walled clear-bottom 96-well plates (Corning Catalog #: 9018) were filled with 180 μl of overnight M9 media (47.8 mM Na_2_HPO_4_, 22 mM KH_2_PO_4_, 8.6 mM NaCl, 18.7 mM NH_4_Cl, 10 mM NaHCO_3_, 0.0025% (w/v) thiamine-HCl, 0.2% (w/v) Casamino Acids, 0.01X Basal Eagle Medium, 0.8% glycerol, 2 mM MgSO_4_, 1 mM CaCl_2_) supplemented with 50 μg/mL kanamycin. We added varying amounts of IPTG to overnight M9 media to achieve the following final concentrations: 0 nM, 50 nM, 10 uM, 50 uM, 500 uM, 1 mM, 5 mM, 10 mM, 50 mM, and 500 mM. Wells were inoculated with 20 μL of each pre-growth culture. Clear lids with condensation rings (Millipore Sigma Catalog #: CLS3931-50EA) were sealed onto plates with parafilm to minimize evaporation. The plate reader (Synergy H1) was pre-heated to 37°C and YFP fluorescence (485 nm/530 nm, gain = 100, measurements/data point = 10), mCherry fluorescence (575 nm/610 nm, measurements/data point = 10), and OD_600_ (measurements/data point = 8) measurements were taken for 16 hours at intervals of 20 minutes (read height = 7 mm, continuous double orbital shaking at frequency = 282 cpm).

### In vivo *data processing*

We normalized each replicate to a control cell line that contains the PAM5087 plasmid but lacks the lac repressor, representing the maximum level of YFP fluorescence possible under each experimental condition. All downstream *in vivo* analyses were conducted at an OD_600_ of 0.53, near mid-log phase. We then fit a Hill model to the YFP vs. [IPTG] data for each genotype. The model we used was:

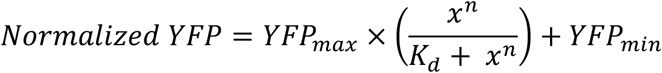

where YFP_max_ corresponds to the maximum YFP fluorescence measured, x is the IPTG concentration, K_d_ is the apparent dissociation constant, n is the Hill coefficient, and YFP_min_ is the minimum YFP fluorescence measured. For genotypes that did not exhibit induction, we instead fit a linear model to the YFP vs. [IPTG] curve. We selected between the Hill and linear model using an AIC test. We calculated the AIC score for the Hill model and the linear models as follows:

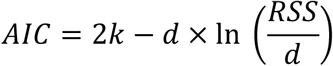

where k is the number of model parameters (k_Hill_ = 3, k_linear_ = 2), d is the number of observations, and RSS is the sum of squared residuals between the model and the experimental data. For each genotype, we selected the model (Hill or linear) with the lower AIC score.

### *Estimating lac repressor concentrations* in vivo

To estimate the concentration of the lac repressor in the *in vivo* experiments, we took an approach similar to Sochor et al ^30^. We cloned the lac repressor-mCherry construct into pBAD with a C-terminal His-tag and purified the protein as above with the untagged constructs. We constructed a BSA calibration curve using SDS-PAGE and gel densitometry. We then used the linear relationship between band density and protein concentration to estimate the concentration of the purified lac repressor-mCherry stock. A calibration curve between protein concentration and mCherry fluorescence was constructed by serially diluting the protein concentration stock (on the same day as the protein concentration vs band density calibration curve) and measuring mCherry fluorescence (excitation: 575 nm, mission: 610 nm) in 96-well black-walled clear-bottom plates (Corning Catalog #: 9018). The linear relationship between concentration and emission intensity at 610 nm was used to convert mCherry signal in all *in vivo* experiments to protein concentration per well (Fig S16).

We converted the concentration of lac repressor per well to an intracellular protein concentration following an approach similar to Sochor et al ^30^. First, we determined the number of cells per well using OD_600_. We measured the OD_600_ for serial dilutions of BLIM cells in 96-well black-walled clear-bottom plates (Corning Catalog #: 9018). We then plated each aliquot on LB agar supplemented with 50 μg/mL kanamycin. After an overnight growth, we counted the number of colonies per plate. We then quantified the relationship between OD_600_ on the plate reader and number of cells by fitting a linear model to the data. We used this calibration curve to calculate the number of cells per well in all *in vivo* experiments.

We determined the total intracellular volume of all cells on the plate by multiplying the number of cells by the volume of an *E. coli* cell (1 × 10^−15^ L)^30,77^. The intracellular protein concentration was then calculated by dividing the concentration of Lac-mCherry per well by the intracellular volume per well. This represents the total concentration of intracellular protein available to bind the O1 operator sequence.

### Protein expression and purification

For all *in vitro* experiments, we used the tetrameric wildtype lac repressor gene in the plasmid phg165c (addgene plasmid #90058) ^44^, obtained courtesy of Prof. Liskin Swint-Kruse. The naturally occurring T109 isoform was used as the wildtype genetic background for all measurements. We used the Quikchange lightning mutagenesis kit to introduce the following three mutations in the lac repressor protein in all possible combinations: M42I, H74A, and K84L. All eight lac repressor mutants were cloned into the pBAD vector with a C-terminal His-tag using a sequence and ligation independent cloning (SLIC) protocol. His-tagged lac repressor pBAD constructs were transformed into ccdB Survival T1R cells for expression.

For protein expression, we inoculated 1.5 L 2xYT (24 g/L Tryptone, 15 g/L Yeast Extract, 7.5 g/L NaCl) supplemented with 50 μg/mL kanamycin with a 1:100 dilution of overnight cultures grew at 37°C, 250 RPM until an OD_600_ between 0.4-0.6 was reached. Protein expression was induced by addition of 0.2% (v/v) L-arabinose and 0.2% glycerol. Cultures were grown for three hours at 37°C, 250 RPM. Cells were harvested by centrifugation at 3000 RPM, 4°C. Cell pellets were resuspended in HisA buffer (46.6 mM Na_2_HPO_4_, 3.4 mM NaH_2_PO_4_, 500 mM NaCl, 20 mM imidazole, 2.5% glycerol, pH 8.0) supplemented with 17.5 U lysozyme/g pellet and 17.5 U DNaseI/g pellet (ThermoFisher, 90082 and 900823). Cells were lysed using sonication with a 30% duty cycle and 55% amplitude in an ice slurry. Lysate was loaded onto two connected 5 mL HisTrap HP columns (Sigma, GE17-5248-02). Protein was purified using an ion exchange gradient protocol by washing with 30 mL HisA buffer and eluting using a 70 mL gradient to 100% HisB (46.6 mM Na_2_HPO_4_, 3.4 mM NaH_2_PO_4_, 500 mM NaCl, 500 mM imidazole, 2.5% glycerol, pH 8.0) on an AKTA Prime FPLC at 4°C. Fractions containing eluted protein were pooled and dialyzed into Binding buffer (42.24 mM Na_2_HPO_4_, 7.75 mM NaH_2_PO_4_, 150 mM NaCl, 1 mM TCEP, 10% glycerol, pH 7.6) overnight at 4°C. Purified protein stocks were confirmed to be >90% pure by SDSPAGE. Protein stocks were flash frozen as pellets in liquid nitrogen and stored at - 80°C.

### Fluorescence polarization measurements

Thawed protein pellets were buffer exchanged into fresh 1X Binding Buffer three times at 15°C, 5000 RPM (42.24 mM Na_2_HPO_4_, 7.75 mM NaH_2_PO_4_, 150 mM NaCl, 1 mM TCEP, 10% glycerol, pH 7.6) using 3 kDa Microsep centrifugal filters (Pall Corporation, #MCP004C41). Scatter-corrected protein concentrations were calculated using A_280_, A_320_, and A_340_ measurements. Lac repressor titrations were prepared in triplicate technical replicates for each biological replicate in black 96-well plates (ThermoScientific catalog #265301). IPTG in 1X Binding Buffer was added to each well to achieve one of the following final concentrations: 0 mM, 0.1 mM, 0.3 mM, or 1 mM. An 18-mer oligo tagged at the 5’ end with fluorescein, FI-Oid (5’-[FI]ATTGTGAGCGCTCACAAT-3’) was ordered HPLC purified from Eurofins MWG and resuspended to 100 μM in 1X TE (pH 8.0) ^78,79^. Resuspended FI-Oid was annealed at a final concentration of 10 μM dsDNA using a thermocycler as follows: 1) 1 cycle at 95°C, 5 minutes 2) 70 cycles at 95°C (-1°C/cycle), 1 minute/cycle. Annealed, double-stranded FI-Oid was then diluted to 100 nM in 1X Binding Buffer and added to each well to achieve a final concentration of 10 nM. Plates were kept covered to protect the fluorescent probe from light and degassed in a temperature-controlled centrifuge pre-equilibrated to 29°C at 300 rcf for 15 minutes. The plate reader (Spectramax i3 equipped with a fluorescence polarization detection cartridge) was pre-equilibrated to 29°C for all experiments. After degassing, samples were shaken in the temperature-equilibrated plate reader (medium orbital) for 15 minutes to ensure the binding reaction reached equilibrium. Fluorescence polarization measurements were taken at an excitation wavelength of 485 nm and an emission wavelength of 535 nm, 6 flashes per read, and 500 ms integration time. Each plate contained four blank wells containing only buffer plus 10 nM FI-Oid and either 0, 0.1, 0.3, or 1 mM IPTG. We calculated a ΔmP value, corresponding to the change in polarization upon the addition of protein, by subtracting the blank from each polarization measurement. We fit the ΔmP values with a single-site binding model to obtain fractional saturation of operator as a function of protein concentration under each IPTG condition.

### Lac repressor ensemble modeling

To extract information about the conformations populated by the lac repressor, we used a previously validated thermodynamic model of the lac repressor ensemble^30^. This model describes the lac repressor with two conformations—high (H) and low (L) DNA affinity—each of which can form a complex with IPTG (I) or DNA (D). Because the lac repressor behaves as a dimer, the repressor dimer can populate states with no effector bound, one effector bound, or two effectors bound. A single dimer binds to a single DNA molecule, meaning each lac repressor is bound to DNA bound or free states. The model assumes the receptor remains a dimer under all conditions. Overall, this model has 12 possible species and five equilibrium constants.

We implemented this model as a function that returns the concentrations of all relevant species given the values of the thermodynamic parameters and total species concentrations of effector, DNA, and lac repressor. This function encodes the thermodynamic relationships derived by Sochor et al. and enforces mass-balance relationships (e.g. the concentrations of the free and bound DNA species must sum to the total DNA concentration). Internally, this function guesses values for free effector and then iterates to self-consistency between the thermodynamic and mass-balance relationships. The software implementing this method—written in a combination of the C and Python programming languages— is available online at https://github.com/harmslab/lacmwc.

To estimate the values of the thermodynamic parameters consistent with our binding and induction data, we used a Bayesian Markov-Chain Monte Carlo (MCMC) strategy to sample over parameter combinations. We analyzed all experimental conditions simultaneously for each genotype, globally estimating the thermodynamic parameters for each genotype. We used the following likelihood function:

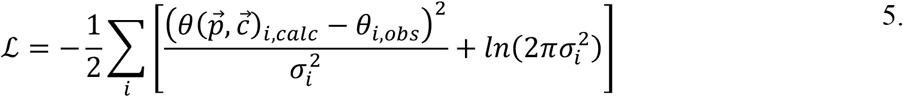

where θ_i,obs_ and σ_i_ are the mean and standard deviation of the measured fractional saturation at condition *i*; θ_i,calc_ is the calculated fractional saturation under these conditions given a vector of parameters and total concentrations. For numerical stability, we did MCMC sampling using the natural logs of the equilibrium constants. We set lower and upper bounds on the sampled ln(K) parameters of -13 and -3.5. These bounds were well below and above the sampled parameter space, but prevented MCMC moves that led to numerical overflows. Within these bounds, we used uninformed priors. We generated 100,000 MCMC samples, discarding the first 10,000 as burn in. We checked for convergence by comparing results from multiple independent runs. We propagated our Bayesian MCMC samples forward through all downstream population and epistasis analyses, leading to the “clouds” of fit lines in all estimates. We used the emcee 3.1.0^80^ and the “likelihood” python libraries (https://github.com/harmslab/likelihood) for these calculations. We implemented our model in Python 3.9 extended with numpy 1.21.1^81^, pandas 1.3.1^82^, and scipy 1.6.2^83^.

We analyzed five variants of the model that differed in the number of free fit parameters (Table S2, Fig S9-S13). We selected between these models using an Akaike Information Criterion (AIC) strategy which penalizes models with excess fit parameters, ultimately find that the “LacMWC3i” model balanced explaining the data against overfitting. The final model had an AIC weight 10^64^ times higher than the next best model (Table S2).

### Far-UV CD Spectroscopy

Far-UV circular dichroism spectra (200–250 nm) were collected on a Model J-815 CD spectrometer (Jasco) with a 1 mm quartz cell (Starna Cells, Inc.) at 25°C. We prepared samples for the untagged and mCherry-tagged wildtype lac repressor constructs at a concentration of 5 μM in 1X Binding Buffer. We collected three scans and averaged to reduce noise. A buffer blank was subtracted from each spectrum and the raw ellipticity was converted to mean molar ellipticity using protein concentrations and the residue length per construct.

## References

(1) Adams, R. M.; Kinney, J. B.; Walczak, A. M.; Mora, T. Epistasis in a Fitness Landscape Defined by Antibody-Antigen Binding Free Energy. Cell Syst. 2019, 8 (1), 86-93.e3. https://doi.org/10.1016/j.cels.2018.12.004.

(2) Anderson, D. W.; McKeown, A. N.; Thornton, J. W. Intermolecular Epistasis Shaped the Function and Evolution of an Ancient Transcription Factor and Its DNA Binding Sites. eLife 2015, 4, e07864. https://doi.org/10.7554/eLife.07864.

(3) Bridgham, J. T.; Carroll, S. M.; Thornton, J. W. Evolution of Hormone-Receptor Complexity by Molecular Exploitation. Science 2006, 312 (5770), 97–101. https://doi.org/10.1126/science.1123348.

(4) Cordell, H. J. Epistasis: What It Means, What It Doesn’t Mean, and Statistical Methods to Detect It in Humans. Hum. Mol. Genet. 2002, 11 (20), 2463–2468. https://doi.org/10.1093/hmg/11.20.2463.

(5) D’Souza, G.; Waschina, S.; Kaleta, C.; Kost, C. Plasticity and Epistasis Strongly Affect Bacterial Fitness after Losing Multiple Metabolic Genes. Evol. Int. J. Org. Evol. 2015, 69 (5), 1244–1254. https://doi.org/10.1111/evo.12640.

(6) Gong, L. I.; Suchard, M. A.; Bloom, J. D. Stability-Mediated Epistasis Constrains the Evolution of an Influenza Protein. eLife 2013, 2, e00631. https://doi.org/10.7554/eLife.00631.

(7) Kachanovsky, D. E.; Filler, S.; Isaacson, T.; Hirschberg, J. Epistasis in Tomato Color Mutations Involves Regulation of Phytoene Synthase 1 Expression by Cis-Carotenoids. Proc. Natl. Acad. Sci. 2012, 109 (46), 19021–19026. https://doi.org/10.1073/pnas.1214808109.

(8) Lunzer, M.; Golding, G. B.; Dean, A. M. Pervasive Cryptic Epistasis in Molecular Evolution. PLOS Genet. 2010, 6 (10), e1001162. https://doi.org/10.1371/journal.pgen.1001162.

(9) McCandlish, D. M.; Otwinowski, J.; Plotkin, J. B. Detecting Epistasis from an Ensemble of Adapting Populations. Evol. Int. J. Org. Evol. 2015, 69 (9), 2359–2370. https://doi.org/10.1111/evo.12735.

(10) Musso, G.; Costanzo, M.; Huangfu, M.; Smith, A. M.; Paw, J.; Luis, B.-J. S.; Boone, C.; Giaever, G.; Nislow, C.; Emili, A.; Zhang, Z. The Extensive and Condition-Dependent Nature of Epistasis among Whole-Genome Duplicates in Yeast. Genome Res. 2008, 18 (7), 1092–1099. https://doi.org/10.1101/gr.076174.108.

(11) Starr, T. N.; Thornton, J. W. Epistasis in Protein Evolution. Protein Sci. 2016, 25 (7), 1204–1218. https://doi.org/10.1002/pro.2897.

(12) Giger, L.; Caner, S.; Obexer, R.; Kast, P.; Baker, D.; Ban, N.; Hilvert, D. Evolution of a Designed Retro-Aldolase Leads to Complete Active Site Remodeling. Nat. Chem. Biol. 2013, 9 (8), 494–498. https://doi.org/10.1038/nchembio.1276.

(13) Miton, C. M.; Tokuriki, N. How Mutational Epistasis Impairs Predictability in Protein Evolution and Design. Protein Sci. 2016, 25 (7), 1260–1272. https://doi.org/10.1002/pro.2876.

(14) Sailer, Z. R.; Harms, M. J. High-Order Epistasis Shapes Evolutionary Trajectories. PLOS Comput. Biol. 2017, 13 (5), e1005541. https://doi.org/10.1371/journal.pcbi.1005541.

(15) Sailer, Z. R.; Harms, M. J. Molecular Ensembles Make Evolution Unpredictable. Proc. Natl. Acad. Sci. 2017, 114 (45), 11938–11943. https://doi.org/10.1073/pnas.1711927114.

(16) Sykora, J.; Brezovsky, J.; Koudelakova, T.; Lahoda, M.; Fortova, A.; Chernovets, T.; Chaloupkova, R.; Stepankova, V.; Prokop, Z.; Smatanova, I. K.; Hof, M.; Damborsky, J. Dynamics and Hydration Explain Failed Functional Transformation in Dehalogenase Design. Nat. Chem. Biol. 2014, 10 (6), 428–430. https://doi.org/10.1038/nchembio.1502.

(17) Weinreich, D. M.; Lan, Y.; Wylie, C. S.; Heckendorn, R. B. Should Evolutionary Geneticists Worry about Higher-Order Epistasis? Curr. Opin. Genet. Dev. 2013, 23 (6), 700–707. https://doi.org/10.1016/j.gde.2013.10.007.

(18) Morrison, A. J.; Wonderlick, D. R.; Harms, M. J. Ensemble Epistasis: Thermodynamic Origins of Nonadditivity between Mutations. Genetics 2021, No. iyab105. https://doi.org/10.1093/genetics/iyab105.

(19) Tsai, C.-J.; Ma, B.; Nussinov, R. Folding and Binding Cascades: Shifts in Energy Landscapes. Proc. Natl. Acad. Sci. 1999, 96 (18), 9970–9972. https://doi.org/10.1073/pnas.96.18.9970.

(20) Tsai, C.-J.; Nussinov, R. A Unified View of “How Allostery Works.” PLOS Comput. Biol. 2014, 10 (2), e1003394. https://doi.org/10.1371/journal.pcbi.1003394.

(21) Smock, R. G.; Gierasch, L. M. Sending Signals Dynamically. Science 2009, 324 (5924), 198–203. https://doi.org/10.1126/science.1169377.

(22) Boehr, D. D.; McElheny, D.; Dyson, H. J.; Wright, P. E. The Dynamic Energy Landscape of Dihydrofolate Reductase Catalysis. Science 2006, 313 (5793), 1638. https://doi.org/10.1126/science.1130258.

(23) del Sol, A.; Tsai, C.-J.; Ma, B.; Nussinov, R. The Origin of Allosteric Functional Modulation: Multiple Pre-Existing Pathways. Structure 2009, 17 (8), 1042–1050. https://doi.org/10.1016/j.str.2009.06.008.

(24) Ma, B.; Nussinov, R. Enzyme Dynamics Point to Stepwise Conformational Selection in Catalysis. Nanotechnol. MiniaturizationMechanisms 2010, 14 (5), 652–659. https://doi.org/10.1016/j.cbpa.2010.08.012.

(25) Motlagh, H. N.; Wrabl, J. O.; Li, J.; Hilser, V. J. The Ensemble Nature of Allostery. Nature 2014, 508 (7496), 331–339. https://doi.org/10.1038/nature13001.

(26) James, L. C.; Roversi, P.; Tawfik, D. S. Antibody Multispecificity Mediated by Conformational Diversity. Science 2003, 299 (5611), 1362. https://doi.org/10.1126/science.1079731.

(27) Yogurtcu, O. N.; Bora Erdemli, S.; Nussinov, R.; Turkay, M.; Keskin, O. Restricted Mobility of Conserved Residues in Protein-Protein Interfaces in Molecular Simulations. Biophys. J. 2008, 94 (9), 3475–3485. https://doi.org/10.1529/biophysj.107.114835.

(28) Hilser, V. J. An Ensemble View of Allostery. Science 2010, 327 (5966), 653–654. https://doi.org/10.1126/science.1186121.

(29) Bell, C. E.; Lewis, M. A Closer View of the Conformation of the Lac Repressor Bound to Operator. Nat. Struct. Biol. 2000, 7 (3), 209–214. https://doi.org/10.1038/73317.

(30) Sochor, M. A. In Vitro Transcription Accurately Predicts Lac Repressor Phenotype in Vivo in Escherichia Coli. PeerJ 2014, 2, e498. https://doi.org/10.7717/peerj.498.

(31) Chuprina, V. P.; Rullmann, J. A. C.; Lamerichs, R. M. J. N.; van Boom, J. H.; Boelens, R.; Kaptein, R. Structure of the Complex of Lac Repressor Headpiece and an 11 Base-Pair Half-Operator Determined by Nuclear Magnetic Resonance Spectroscopy and Restrained Molecular Dynamics. J. Mol. Biol. 1993, 234 (2), 446–462. https://doi.org/10.1006/jmbi.1993.1598.

(32) Friedman, A. M.; Fischmann, T. O.; Steitz, T. A. Crystal Structure of Lac Repressor Core Tetramer and Its Implications for DNA Looping. Science 1995, 268 (5218), 1721–1727. https://doi.org/10.1126/science.7792597.

(33) Gilbert, W.; Müller-Hill, B. Isolation of the Lac Repressor. Proc. Natl. Acad. Sci. 1966, 56 (6), 1891–1898. https://doi.org/10.1073/pnas.56.6.1891.

(34) Jacob, F.; Monod, J. Genetic Regulatory Mechanisms in the Synthesis of Proteins. J. Mol. Biol. 1961, 3 (3), 318–356. https://doi.org/10.1016/S0022-2836(61)80072-7.

(35) Lewis, M. The Lac Repressor. C. R. Biol. 2005, 328 (6), 521–548. https://doi.org/10.1016/j.crvi.2005.04.004.

(36) Lewis, M.; Chang, G.; Horton, N. C.; Kercher, M. A.; Pace, H. C.; Schumacher, M. A.; Brennan, R. G.; Lu, P. Crystal Structure of the Lactose Operon Repressor and Its Complexes with DNA and Inducer. Science 1996, 271 (5253), 1247–1254. https://doi.org/10.1126/science.271.5253.1247.

(37) Monod, J.; Wyman, J.; Changeux, J.-P. On the Nature of Allosteric Transitions: A Plausible Model. J. Mol. Biol. 1965, 12 (1), 88–118. https://doi.org/10.1016/S0022-2836(65)80285-6.

(38) Müller-Hill, B.; Rickenberg, H. V.; Wallenfels, K. Specificity of the Induction of the Enzymes of the Lac Operon in Escherichia Coli. J. Mol. Biol. 1964, 10 (2), 303–318. https://doi.org/10.1016/S0022-2836(64)80049-8.

(39) Perez, P. J.; Clauvelin, N.; Tam, G.; Olson, W. K. Conformational Changes in the Lac Repressor Protein Effect DNA Loop Energetics and Topology. Biophys. J. 2014, 106 (2), 71a. https://doi.org/10.1016/j.bpj.2013.11.467.

(40) Slijper, M.; Bonvin, A. M. J. J.; Boelens, R.; Kaptein, R. Refined Structure of Lac Repressor Headpiece (1-56) Determined by Relaxation Matrix Calculations from 2D and 3D NOE Data: Change of Tertiary Structure upon Binding to ThelacOperator. J. Mol. Biol. 1996, 259 (4), 761–773. https://doi.org/10.1006/jmbi.1996.0356.

(41) Taraban, M.; Zhan, H.; Whitten, A. E.; Langley, D. B.; Matthews, K. S.; Swint-Kruse, L.; Trewhella, J. Ligand-Induced Conformational Changes and Conformational Dynamics in the Solution Structure of the Lactose Repressor Protein. J. Mol. Biol. 2008, 376 (2), 466–481. https://doi.org/10.1016/j.jmb.2007.11.067.

(42) Vos, M. G. J. de; Poelwijk, F. J.; Battich, N.; Ndika, J. D. T.; Tans, S. J. Environmental Dependence of Genetic Constraint. PLOS Genet. 2013, 9 (6), e1003580. https://doi.org/10.1371/journal.pgen.1003580.

(43) Barry, J. K.; Matthews, K. S. Substitutions at Histidine 74 and Aspartate 278 Alter Ligand Binding and Allostery in Lactose Repressor Protein. Biochemistry 1999, 38 (12), 3579–3590. https://doi.org/10.1021/bi982577n.

(44) Meinhardt, S.; Manley, M. W., Jr; Becker, N. A.; Hessman, J. A.; Maher, L. J., III; Swint-Kruse, L. Novel Insights from Hybrid LacI/GalR Proteins: Family-Wide Functional Attributes and Biologically Significant Variation in Transcription Repression. Nucleic Acids Res. 2012, 40 (21), 11139–11154. https://doi.org/10.1093/nar/gks806.

(45) Swint-Kruse, L.; Elam, C. R.; Lin, J. W.; Wycuff, D. R.; Matthews, K. S. Plasticity of Quaternary Structure: Twenty-Two Ways to Form a LacI Dimer. Protein Sci. Publ. Protein Soc. 2001, 10 (2), 262–276.

(46) Swint-Kruse, L.; Zhan, H.; Matthews, K. S. Integrated Insights from Simulation, Experiment, and Mutational Analysis Yield New Details of LacI Function. Biochemistry 2005, 44 (33), 11201–11213. https://doi.org/10.1021/bi050404+.

(47) Bell, C. E.; Barry, J.; Matthews, K. S.; Lewis, M. Structure of a Variant of Lac Repressor with Increased Thermostability and Decreased Affinity for Operator11Edited by D. Rees. J. Mol. Biol. 2001, 313 (1), 99–109. https://doi.org/10.1006/jmbi.2001.5041.

(48) Flynn, T. C.; Swint-Kruse, L.; Kong, Y.; Booth, C.; Matthews, K. S.; Ma, J. Allosteric Transition Pathways in the Lactose Repressor Protein Core Domains: Asymmetric Motions in a Homodimer. Protein Sci. Publ. Protein Soc. 2003, 12 (11), 2523–2541.

(49) Sailer, Z. R.; Harms, M. J. Detecting High-Order Epistasis in Nonlinear Genotype-Phenotype Maps. Genetics 2017, 205 (3), 1079–1088. https://doi.org/10.1534/genetics.116.195214.

(50) Taute, K. M.; Gude, S.; Nghe, P.; Tans, S. J. Evolutionary Constraints in Variable Environments, from Proteins to Networks. Trends Genet. 2014, 30 (5), 192–198. https://doi.org/10.1016/j.tig.2014.04.003.

(51) Otwinowski, J. Biophysical Inference of Epistasis and the Effects of Mutations on Protein Stability and Function. Mol. Biol. Evol. 2018, 35 (10), 2345–2354. https://doi.org/10.1093/molbev/msy141.

(52) Anderson, D. W.; Baier, F.; Yang, G.; Tokuriki, N. The Adaptive Landscape of a Metallo-Enzyme Is Shaped by Environment-Dependent Epistasis. Nat. Commun. 2021, 12 (1), 3867. https://doi.org/10.1038/s41467-021-23943-x.

(53) Hayden, E. J.; Ferrada, E.; Wagner, A. Cryptic Genetic Variation Promotes Rapid Evolutionary Adaptation in an RNA Enzyme. Nature 2011, 474 (7349), 92–95. https://doi.org/10.1038/nature10083.

(54) Hayden, E. J.; Wagner, A. Environmental Change Exposes Beneficial Epistatic Interactions in a Catalytic RNA. Proc. R. Soc. B Biol. Sci. 2012, 279 (1742), 3418–3425. https://doi.org/10.1098/rspb.2012.0956.

(55) Guerrero, R. F.; Scarpino, S. V.; Rodrigues, J. V.; Hartl, D. L.; Ogbunugafor, C. B. Proteostasis Environment Shapes Higher-Order Epistasis Operating on Antibiotic Resistance. Genetics 2019, 212 (2), 565–575. https://doi.org/10.1534/genetics.119.302138.

(56) Lindsey, H. A.; Gallie, J.; Taylor, S.; Kerr, B. Evolutionary Rescue from Extinction Is Contingent on a Lower Rate of Environmental Change. Nature 2013, 494 (7438), 463–467. https://doi.org/10.1038/nature11879.

(57) Ogbunugafor, C. B.; Wylie, C. S.; Diakite, I.; Weinreich, D. M.; Hartl, D. L. Adaptive Landscape by Environment Interactions Dictate Evolutionary Dynamics in Models of Drug Resistance. PLOS Comput. Biol. 2016, 12 (1), e1004710. https://doi.org/10.1371/journal.pcbi.1004710.

(58) Li, C.; Zhang, J. Multi-Environment Fitness Landscapes of a TRNA Gene. Nat. Ecol. Evol. 2018, 2 (6), 1025–1032. https://doi.org/10.1038/s41559-018-0549-8.

(59) Lagator, M.; Igler, C.; Moreno, A. B.; Guet, C. C.; Bollback, J. P. Epistatic Interactions in the Arabinose Cis-Regulatory Element. Mol. Biol. Evol. 2016, 33 (3), 761–769. https://doi.org/10.1093/molbev/msv269.

(60) Lagator, M.; Paixão, T.; Barton, N. H.; Bollback, J. P.; Guet, C. C. On the Mechanistic Nature of Epistasis in a Canonical Cis-Regulatory Element. eLife 2017, 6, e25192. https://doi.org/10.7554/eLife.25192.

(61) Shultzaberger, R. K.; Malashock, D. S.; Kirsch, J. F.; Eisen, M. B. The Fitness Landscapes of Cis-Acting Binding Sites in Different Promoter and Environmental Contexts. PLOS Genet. 2010, 6 (7), e1001042. https://doi.org/10.1371/journal.pgen.1001042.

(62) Nghe, P.; Kogenaru, M.; Tans, S. J. Sign Epistasis Caused by Hierarchy within Signalling Cascades. Nat. Commun. 2018, 9 (1), 1451. https://doi.org/10.1038/s41467-018-03644-8.

(63) Caudle, S. B.; Miller, C. R.; Rokyta, D. R. Environment Determines Epistatic Patterns for a SsDNA Virus. Genetics 2014, 196 (1), 267–279. https://doi.org/10.1534/genetics.113.158154.

(64) Filteau, M.; Hamel, V.; Pouliot, M.-C.; Gagnon-Arsenault, I.; Dubé, A. K.; Landry, C. R. Evolutionary Rescue by Compensatory Mutations Is Constrained by Genomic and Environmental Backgrounds. Mol. Syst. Biol. 2015, 11 (10), 832. https://doi.org/10.15252/msb.20156444.

(65) Flynn, K. M.; Cooper, T. F.; Moore, F. B.-G.; Cooper, V. S. The Environment Affects Epistatic Interactions to Alter the Topology of an Empirical Fitness Landscape. PLOS Genet. 2013, 9 (4), e1003426. https://doi.org/10.1371/journal.pgen.1003426.

(66) James, L. C.; Tawfik, D. S. Conformational Diversity and Protein Evolution – a 60-Year-Old Hypothesis Revisited. Trends Biochem. Sci. 2003, 28 (7), 361–368. https://doi.org/10.1016/S0968-0004(03)00135-X.

(67) Petrović, D.; Risso, V. A.; Kamerlin, S. C. L.; Sanchez-Ruiz, J. M. Conformational Dynamics and Enzyme Evolution. J. R. Soc. Interface 2018, 15 (144), 20180330. https://doi.org/10.1098/rsif.2018.0330.

(68) Tokuriki, N.; Tawfik, D. S. Protein Dynamism and Evolvability. Science 2009, 324 (5924), 203–207. https://doi.org/10.1126/science.1169375.

(69) Jolly, M. K.; Kulkarni, P.; Weninger, K.; Orban, J.; Levine, H. Phenotypic Plasticity, Bet-Hedging, and Androgen Independence in Prostate Cancer: Role of Non-Genetic Heterogeneity. Front. Oncol. 2018, 8, 50. https://doi.org/10.3389/fonc.2018.00050.

(70) Pigliucci, M.; Murren, C. J.; Schlichting, C. D. Phenotypic Plasticity and Evolution by Genetic Assimilation. J. Exp. Biol. 2006, 209 (12), 2362–2367. https://doi.org/10.1242/jeb.02070.

(71) Javier Zea, D.; Miguel Monzon, A.; Fornasari, M. S.; Marino-Buslje, C.; Parisi, G. Protein Conformational Diversity Correlates with Evolutionary Rate. Mol. Biol. Evol. 2013, 30 (7), 1500–1503. https://doi.org/10.1093/molbev/mst065.

(72) Tóth-Petróczy, Á.; Tawfik, D. S. The Robustness and Innovability of Protein Folds. Curr. Opin. Struct. Biol. 2014, 26, 131–138. https://doi.org/10.1016/j.sbi.2014.06.007.

(73) Cohen, S. E.; Erb, M. L.; Selimkhanov, J.; Dong, G.; Hasty, J.; Pogliano, J.; Golden, S. S. Dynamic Localization of the Cyanobacterial Circadian Clock Proteins. Curr. Biol. 2014, 24 (16), 1836–1844. https://doi.org/10.1016/j.cub.2014.07.036.

(74) Arai, R.; Ueda, H.; Kitayama, A.; Kamiya, N.; Nagamune, T. Design of the Linkers Which Effectively Separate Domains of a Bifunctional Fusion Protein. Protein Eng. Des. Sel. 2001, 14 (8), 529–532. https://doi.org/10.1093/protein/14.8.529.

(75) Lau, I. F.; Filipe, S. R.; Søballe, B.; Økstad, O.-A.; Barre, F.-X.; Sherratt, D. J. Spatial and Temporal Organization of Replicating Escherichia Coli Chromosomes. Mol. Microbiol. 2003, 49 (3), 731–743. https://doi.org/10.1046/j.1365-2958.2003.03640.x.

(76) Wycuff, D. R.; Matthews, K. S. Generation of an AraC-AraBAD Promoter-Regulated T7 Expression System. Anal. Biochem. 2000, 277 (1), 67–73. https://doi.org/10.1006/abio.1999.4385.

(77) Kubitschek, H. E.; Friske, J. A. Determination of Bacterial Cell Volume with the Coulter Counter. J. Bacteriol. 1986, 168 (3), 1466–1467. https://doi.org/10.1128/jb.168.3.1466-1467.1986.

(78) Hoffmann, S. A.; Kruse, S. M.; Arndt, K. M. Long-Range Transcriptional Interference in E. Coli Used to Construct a Dual Positive Selection System for Genetic Switches. Nucleic Acids Res. 2016, 44 (10), e95. https://doi.org/10.1093/nar/gkw125.

(79) Poelwijk, F. J.; de Vos, M. G. J.; Tans, S. J. Tradeoffs and Optimality in the Evolution of Gene Regulation. Cell 2011, 146 (3), 462–470. https://doi.org/10.1016/j.cell.2011.06.035.

(80) Foreman-Mackey, D.; Hogg, D. W.; Lang, D.; Goodman, J. Emcee: The MCMC Hammer. Publ. Astron. Soc. Pac. 2013, 125 (925), 306. https://doi.org/10.1086/670067.

(81) Harris, C. R.; Millman, K. J.; van der Walt, S. J.; Gommers, R.; Virtanen, P.; Cournapeau, D.; Wieser, E.; Taylor, J.; Berg, S.; Smith, N. J.; Kern, R.; Picus, M.; Hoyer, S.; van Kerkwijk, M. H.; Brett, M.; Haldane, A.; del Río, J. F.; Wiebe, M.; Peterson, P.; Gérard-Marchant, P.; Sheppard, K.; Reddy, T.; Weckesser, W.; Abbasi, H.; Gohlke, C.; Oliphant, T. E. Array Programming with NumPy. Nature 2020, 585 (7825), 357–362. https://doi.org/10.1038/s41586-020-2649-2.

(82) Reback, J.; jbrockmendel McKinney, W.; Bossche, J. V. den; Augspurger, T.; Cloud, P.; Hawkins, S.; gfyoung Sinhrks Roeschke, M.; Klein, A.; Petersen, T.; Tratner, J.; She, C.; Ayd, W.; Hoefler, P.; Naveh, S.; Garcia, M.; Schendel, J.; Hayden, A.; Saxton, D.; Shadrach, R.; Gorelli, M. E.; Li, F.; Jancauskas, V.; attack68; McMaster, A.; Battiston, P.; Seabold, S.; Dong, K. Pandas-Dev/Pandas: Pandas 1.3.2; Zenodo, 2021. https://doi.org/10.5281/zenodo.5203279.

(83) Virtanen, P.; Gommers, R.; Oliphant, T. E.; Haberland, M.; Reddy, T.; Cournapeau, D.; Burovski, E.; Peterson, P.; Weckesser, W.; Bright, J.; van der Walt, S. J.; Brett, M.; Wilson, J.; Millman, K. J.; Mayorov, N.; Nelson, A. R. J.; Jones, E.; Kern, R.; Larson, E.; Carey, C. J.; Polat, İ.; Feng, Y.; Moore, E. W.; VanderPlas, J.; Laxalde, D.; Perktold, J.; Cimrman, R.; Henriksen, I.; Quintero, E. A.; Harris, C. R.; Archibald, A. M.; Ribeiro, A. H.; Pedregosa, F.; van Mulbregt, P.; SciPy 1.0 Contributors. SciPy 1.0: Fundamental Algorithms for Scientific Computing in Python. Nat. Methods 2020, 17, 261–272. https://doi.org/10.1038/s41592-019-0686-2.

