## Supplementary material for "An experimental demonstration of ensemble epistasis in the lac repressor": Fig S1-S16, Table S1-S2

### mhk

Right: Binding of repressor to operator DNA *in vitro*, as measured by fluorescence anisotropy. Plot shows fractional saturation of operator versus repressor concentration. Points and error bars indicate the average and standard deviation of at least three biological replicates. Clouds of lines show model parameter sets from a global Bayesian MCMC analysis. The color of each dataset corresponds to the IPTG concentration as indicated on the figure. Note: Each IPTG series included a 0 M repressor sample that is not shown on the plot.

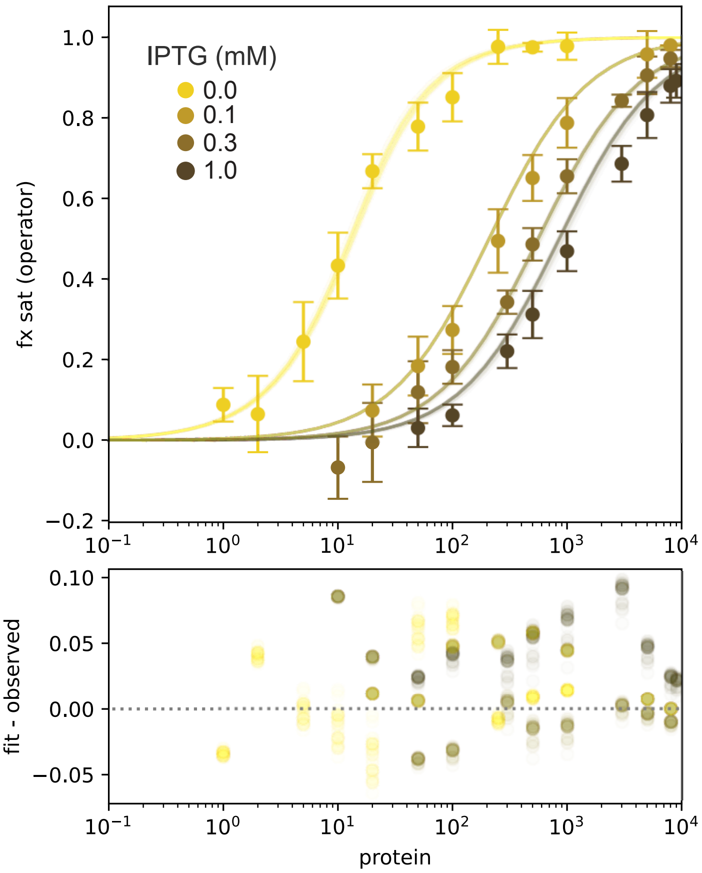


Below: corner plots show marginal distribution of values for each model parameter (panels on diagonal) and correlation of fit parameters (panels off-diagonal) across MCMC samples. Blue lines indicate median values. “s” is a nuisance paramter scaling operator affinity in vitro relative to its value in vivo.

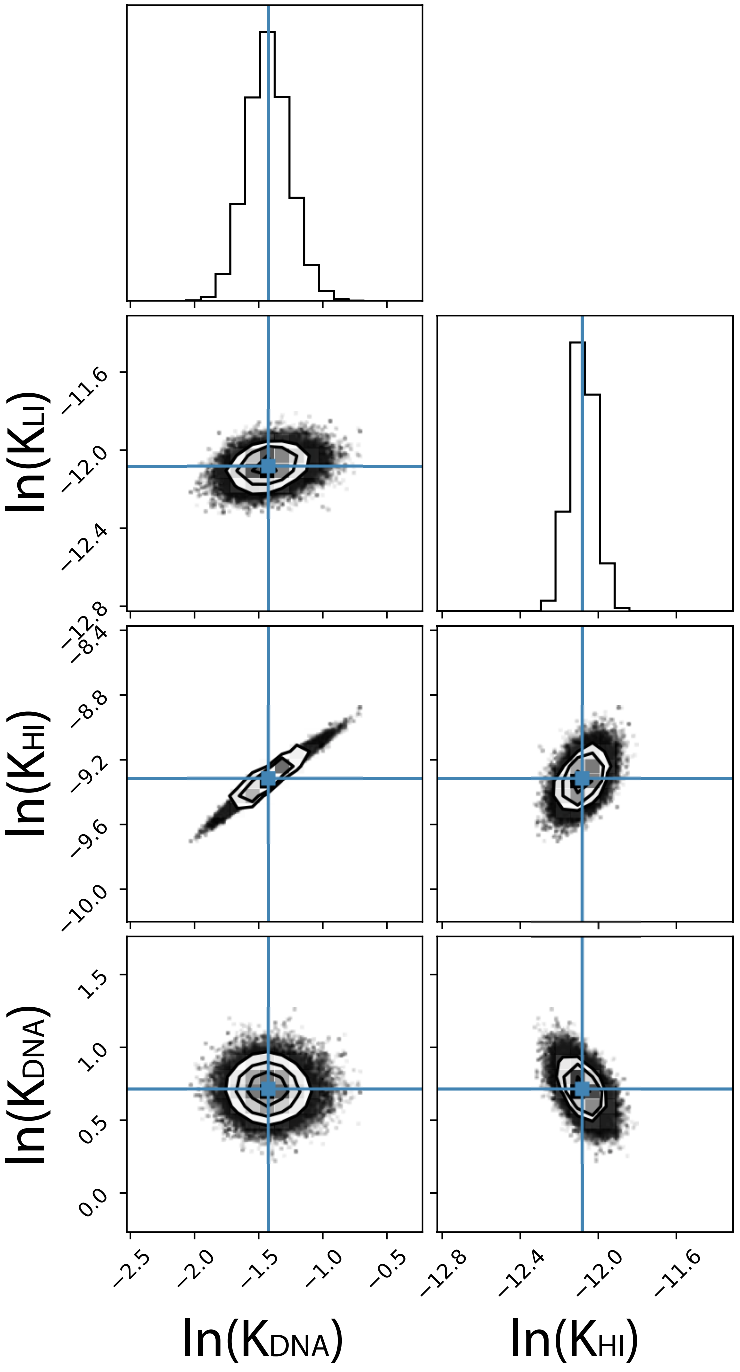


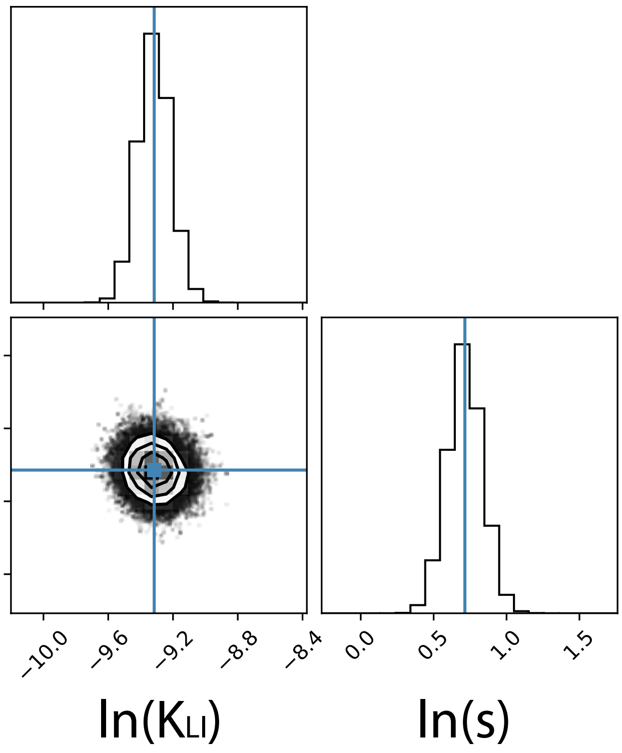

Right: Binding of repressor to operator DNA *in vitro*, as measured by fluorescence anisotropy. Plot shows fractional saturation of operator versus repressor concentration. Points and error bars indicate the average and standard deviation of at least three biological replicates. Clouds of lines show model parameter sets from a global Bayesian MCMC analysis. The color of each dataset corresponds to the IPTG concentration as indicated on the figure. Note: Each IPTG series included a 0 M repressor sample that is not shown on the plot.

Below: corner plots show marginal distribution of values for each model parameter (panels on diagonal) and correlation of fit parameters (panels off-diagonal) across MCMC samples. Blue lines indicate median values. “s” is a nuisance paramter scaling operator affinity in vitro relative to its value in vivo.

### Fig S2

#### lhk

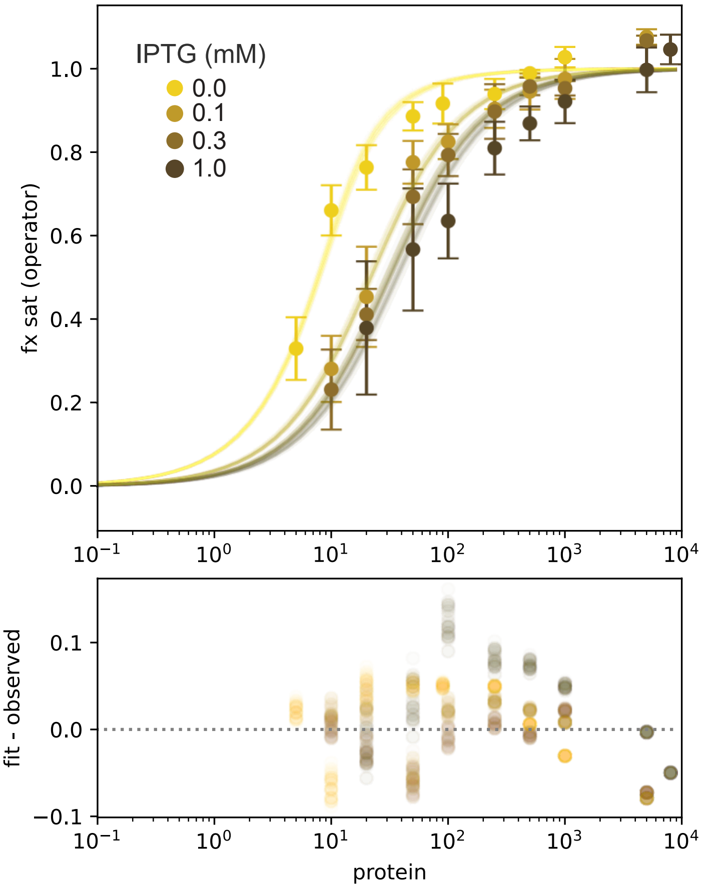


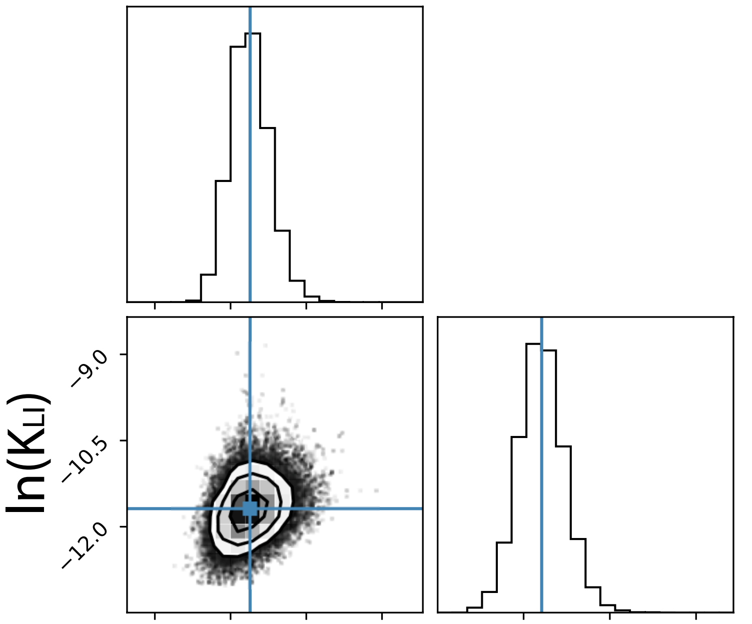


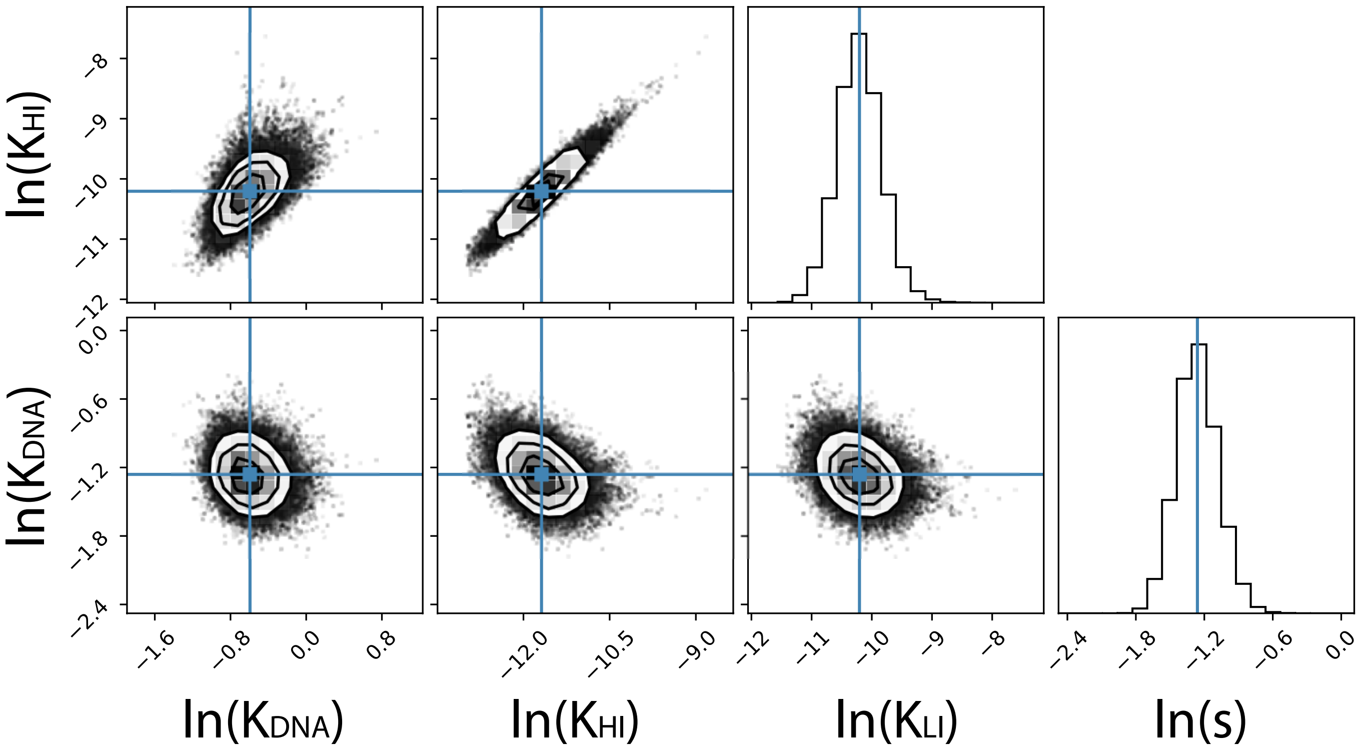

Right: Binding of repressor to operator DNA *in vitro*, as measured by fluorescence anisotropy. Plot shows fractional saturation of operator versus repressor concentration. Points and error bars indicate the average and standard deviation of at least three biological replicates. Clouds of lines show model parameter sets from a global Bayesian MCMC analysis. The color of each dataset corresponds to the IPTG concentration as indicated on the figure. Note: Each IPTG series included a 0 M repressor sample that is not shown on the plot.

Below: corner plots show marginal distribution of values for each model parameter (panels on diagonal) and correlation of fit parameters (panels off-diagonal) across MCMC samples. Blue lines indicate median values.  “s” is a nuisance paramter scaling operator affinity in vitro relative to its value in vivo.

### Fig S3

#### mAk

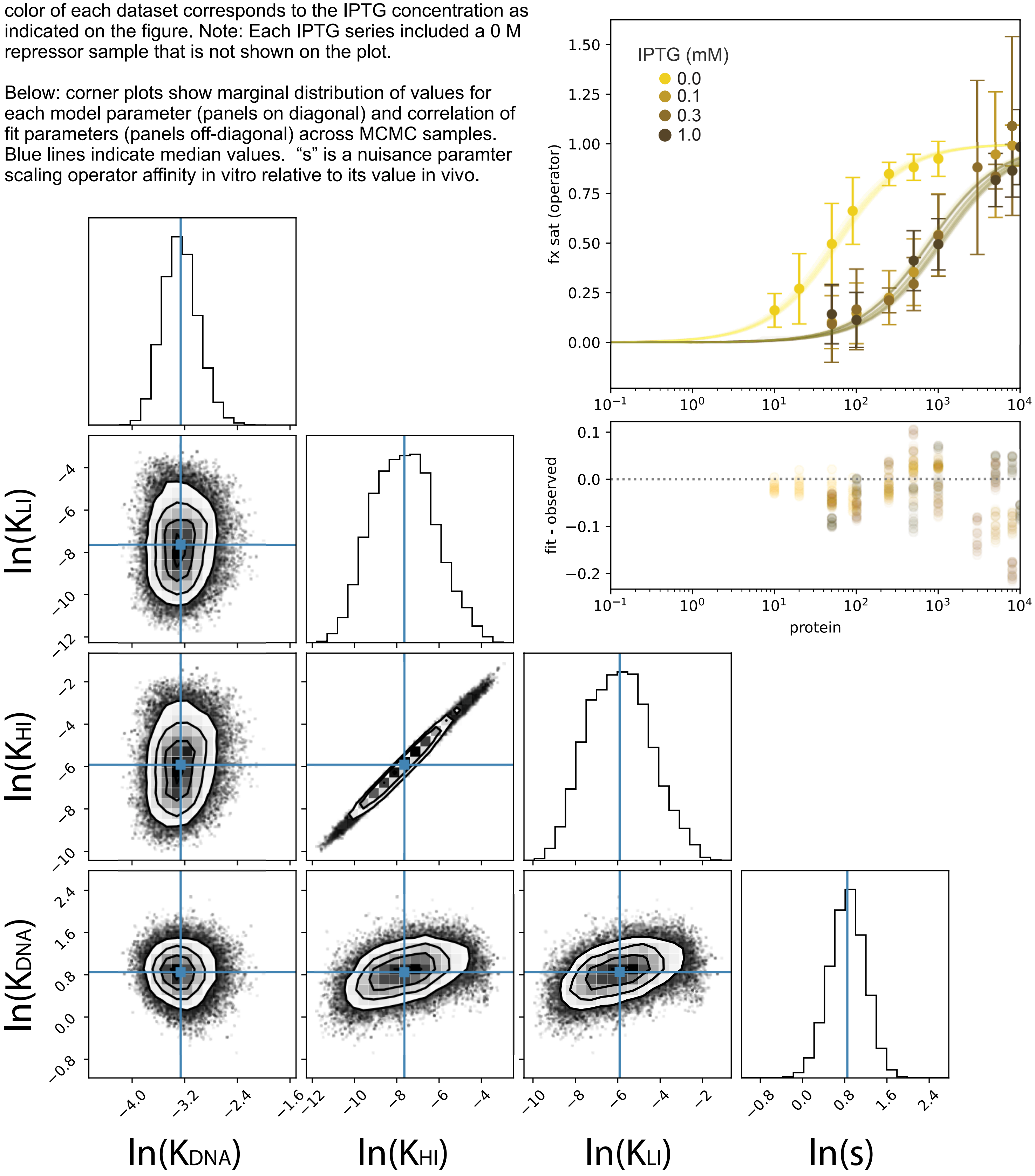

Right: Binding of repressor to operator DNA *in vitro*, as measured by fluorescence anisotropy. Plot shows fractional saturation of operator versus repressor concentration. Points and error bars indicate the average and standard deviation of at least three biological replicates. Clouds of lines show model parameter sets from a global Bayesian MCMC analysis. The color of each dataset corresponds to the IPTG concentration as indicated on the figure. Note: Each IPTG series included a 0 M repressor sample that is not shown on the plot.

Below: corner plots show marginal distribution of values for each model parameter (panels on diagonal) and correlation of fit parameters (panels off-diagonal) across MCMC samples. Blue lines indicate median values. “s” is a nuisance paramter scaling operator affinity in vitro relative to its value in vivo.

### Fig S4

#### mhL

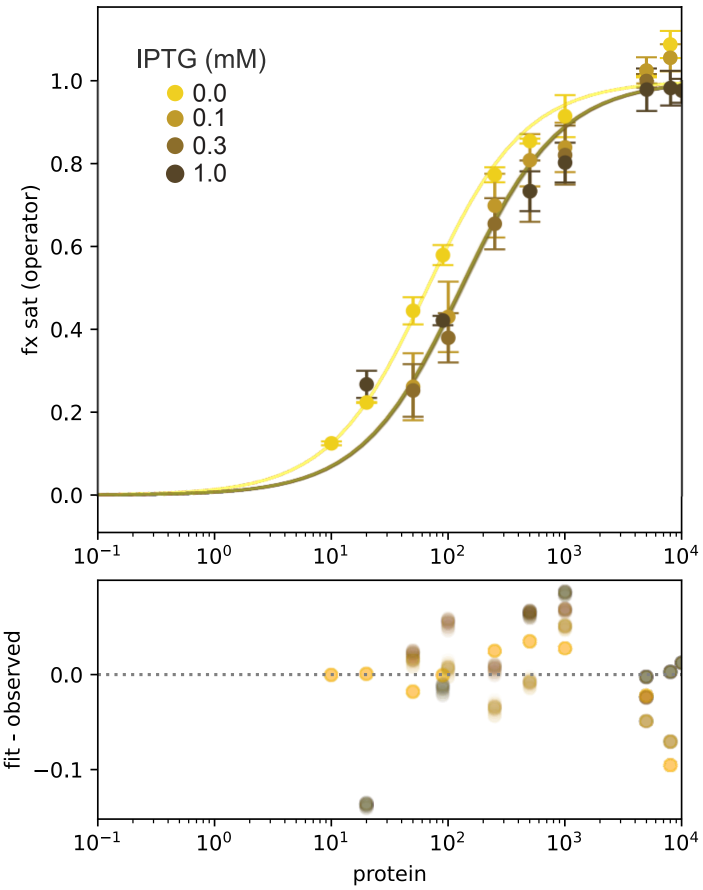


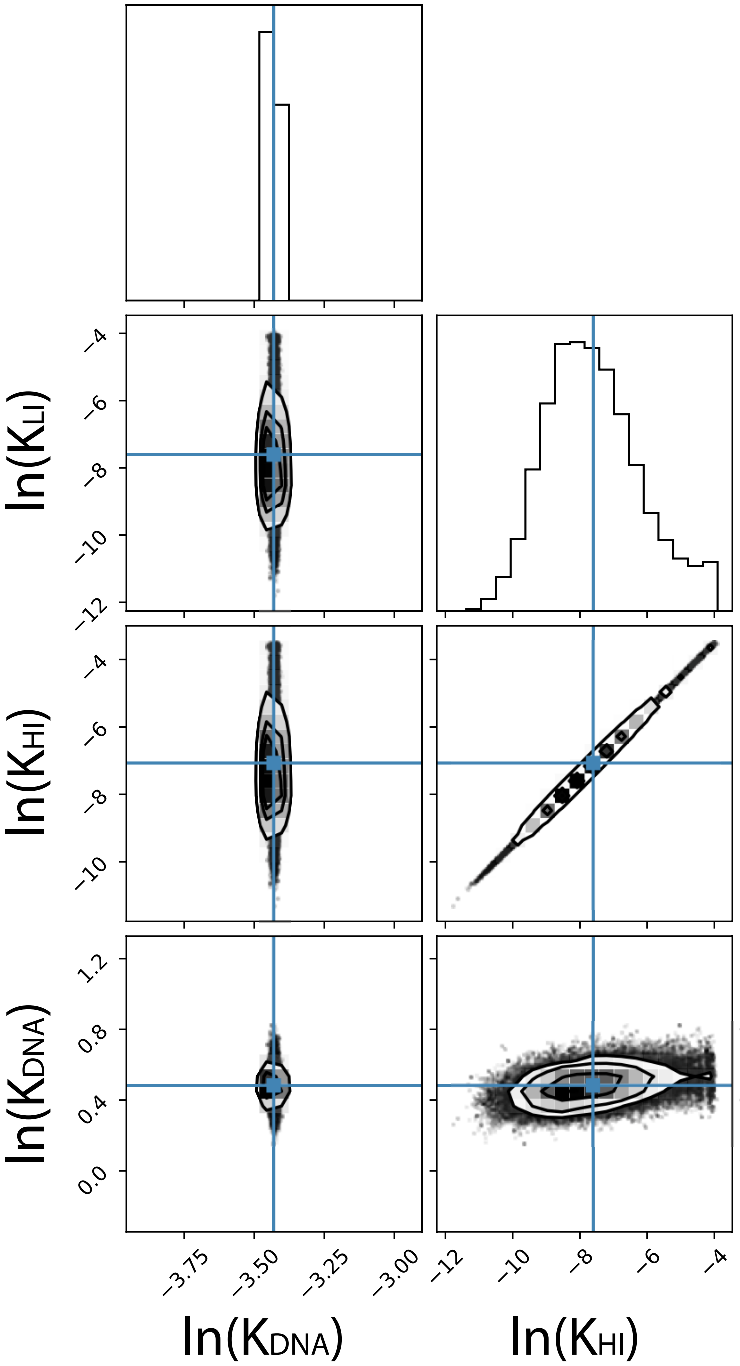
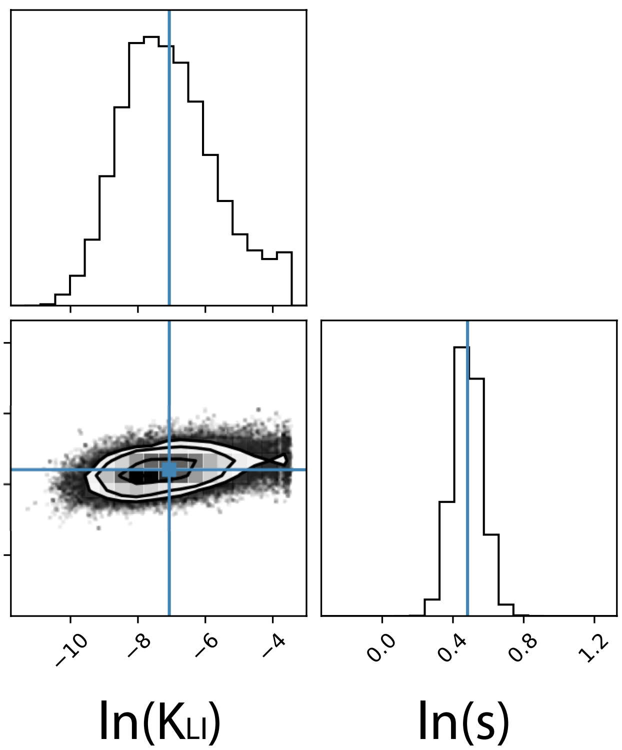

Right: Binding of repressor to operator DNA *in vitro*, as measured by fluorescence anisotropy. Plot shows fractional saturation of operator versus repressor concentration. Points and error bars indicate the average and standard deviation of at least three biological replicates. Clouds of lines show model parameter sets from a global Bayesian MCMC analysis. The color of each dataset corresponds to the IPTG concentration as indicated on the figure. Note: Each IPTG series included a 0 M repressor sample that is not shown on the plot.

Below: corner plots show marginal distribution of values for each model parameter (panels on diagonal) and correlation of fit parameters (panels off-diagonal) across MCMC samples. Blue lines indicate median values. “s” is a nuisance paramter scaling operator affinity in vitro relative to its value in vivo.

Fig S5

### IAk

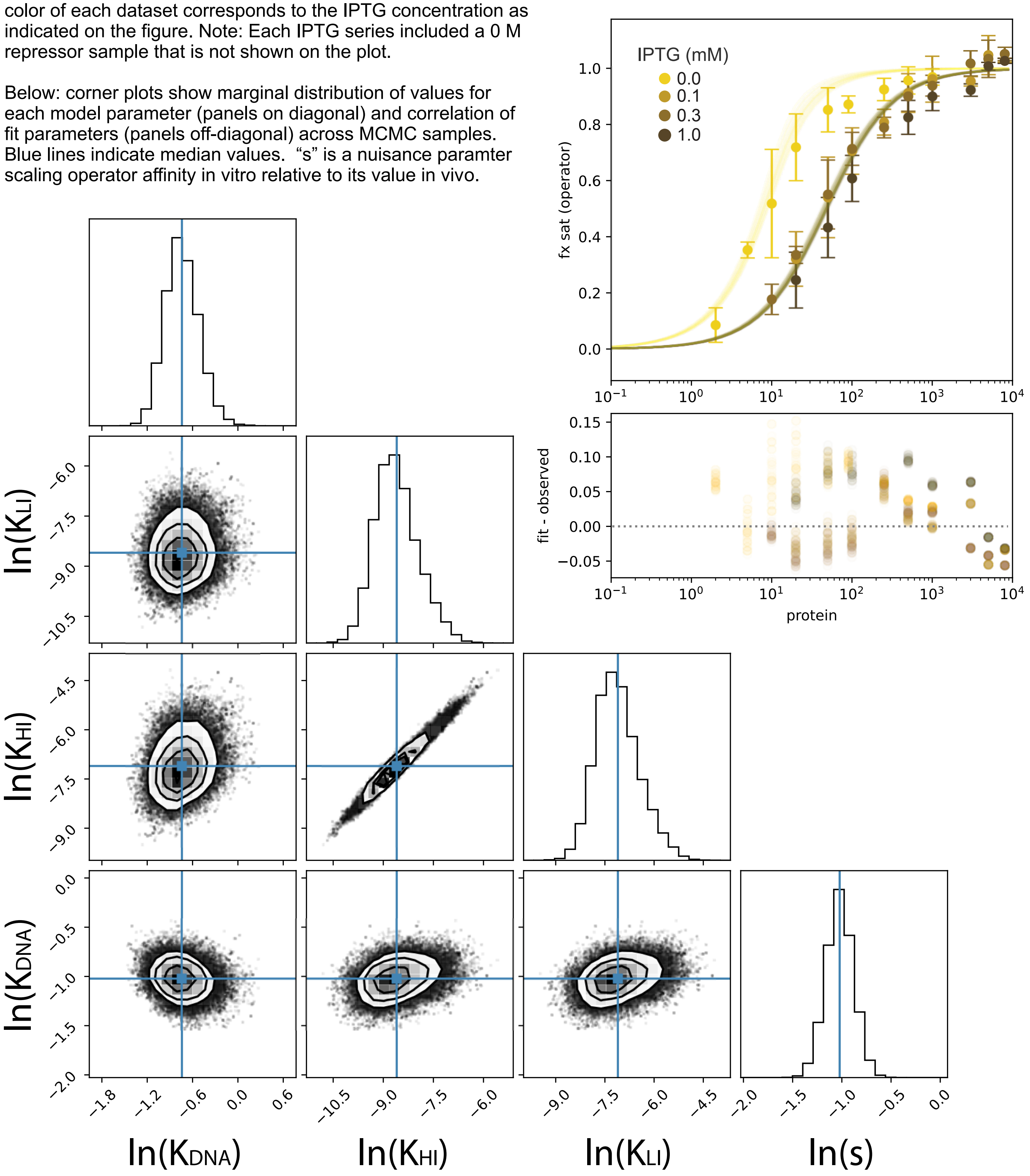

Right: Binding of repressor to operator DNA *in vitro*, as measured by fluorescence anisotropy. Plot shows fractional saturation of operator versus repressor concentration. Points and error bars indicate the average and standard deviation of at least three biological replicates. Clouds of lines show model parameter sets from a global Bayesian MCMC analysis. The color of each dataset corresponds to the IPTG concentration as indicated on the figure. Note: Each IPTG series included a 0 M repressor sample that is not shown on the plot.

Below: corner plots show marginal distribution of values for each model parameter (panels on diagonal) and correlation of fit parameters (panels off-diagonal) across MCMC samples. Blue lines indicate median values. “s” is a nuisance paramter scaling operator affinity in vitro relative to its value in vivo.

### Fig S6

#### IhL

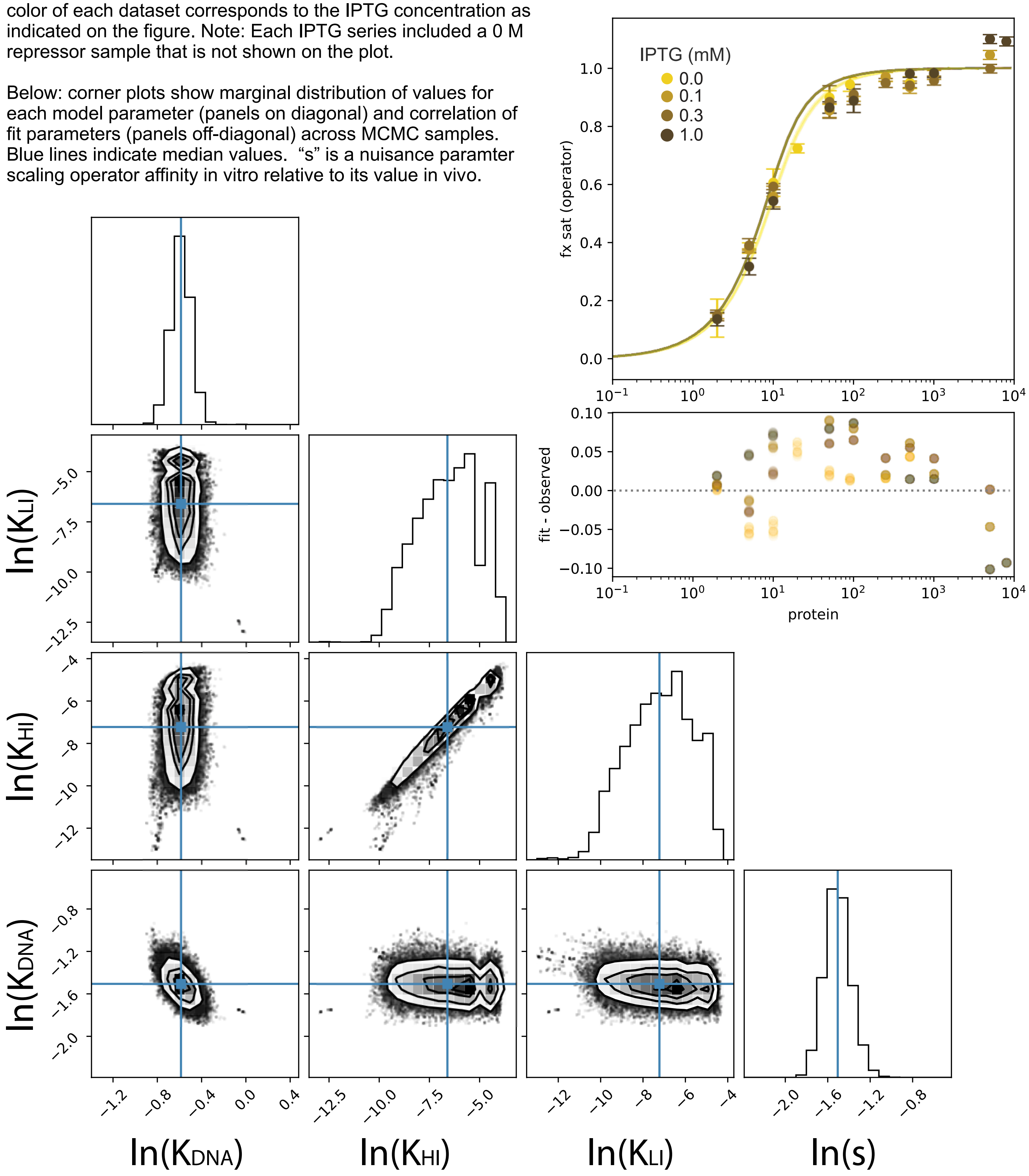

### Fig S7

### mAL

Right: Binding of repressor to operator DNA *in vitro*, as measured by fluorescence anisotropy. Plot shows fractional saturation of operator versus repressor concentration. Points and error bars indicate the average and standard deviation of at least three biological replicates. Clouds of lines show model parameter sets from a global Bayesian MCMC analysis. The color of each dataset corresponds to the IPTG concentration as indicated on the figure. Note: Each IPTG series included a 0 M repressor sample that is not shown on the plot.

Below: corner plots show marginal distribution of values for each model parameter (panels on diagonal) and correlation of fit parameters (panels off-diagonal) across MCMC samples. Blue lines indicate median values. “s” is a nuisance paramter scaling operator affinity in vitro relative to its value in vivo.

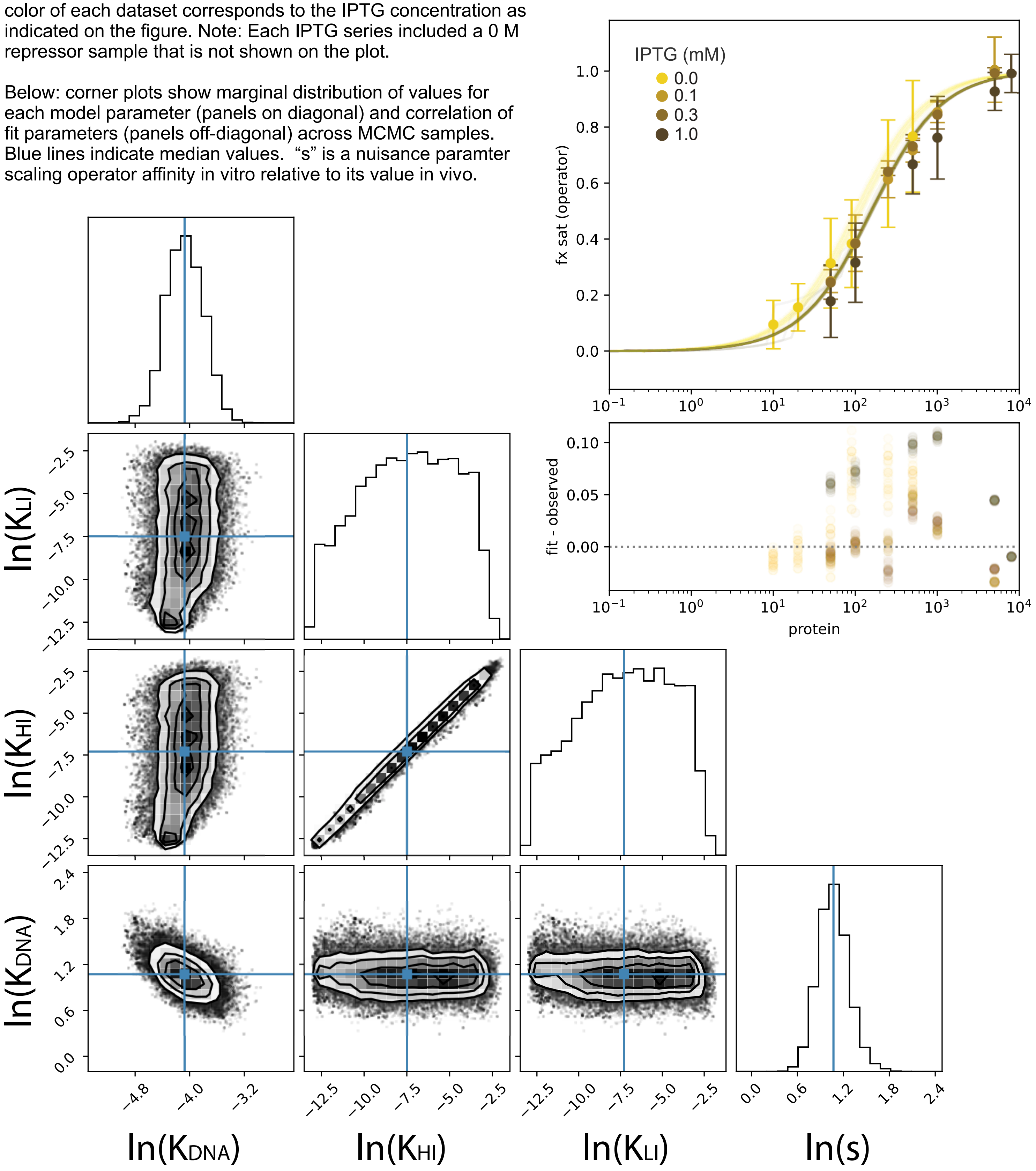

Fig S8

### IAL

Right: Binding of repressor to operator DNA *in vitro*, as measured by fluorescence anisotropy. Plot shows fractional saturation of operator versus repressor concentration. Points and error bars indicate the average and standard deviation of at least three biological replicates. Clouds of lines show model parameter sets from a global Bayesian MCMC analysis. The color of each dataset corresponds to the IPTG concentration as indicated on the figure. Note: Each IPTG series included a 0 M repressor sample that is not shown on the plot.

Below: corner plots show marginal distribution of values for each model parameter (panels on diagonal) and correlation of fit parameters (panels off-diagonal) across MCMC samples. Blue lines indicate median values. “s” is a nuisance paramter scaling operator affinity in vitro relative to its value in vivo.

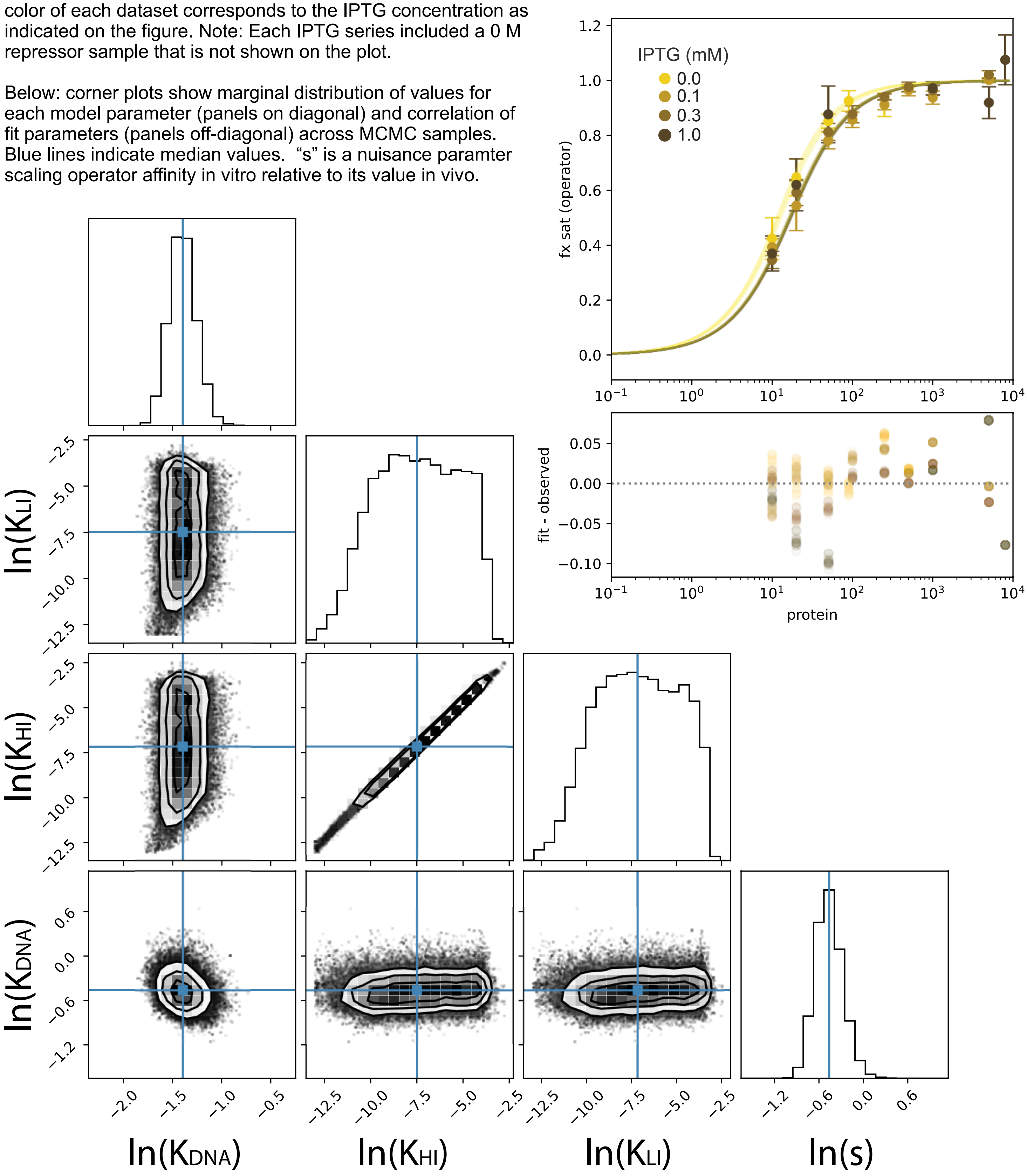

**Fig S9: Corner plot for MCMC samples of parameters for the LacMWC5 model against the binding data for the mhk genotype.** Parameters: $\ln(K_{H\cdot I})$, $\ln(K_{L\cdot I})$, $\ln(K_{H\cdot DNA})$, $\ln(K_{L\cdot DNA})$, $\ln(K_{HL})$. Plots show marginal distribution of values for each model parameter (panels on diagonal) and correlation of fit parameters (panels off-diagonal) across MCMC samples. Blue lines indicate median values. This is the model used in the analyses in this manuscript.

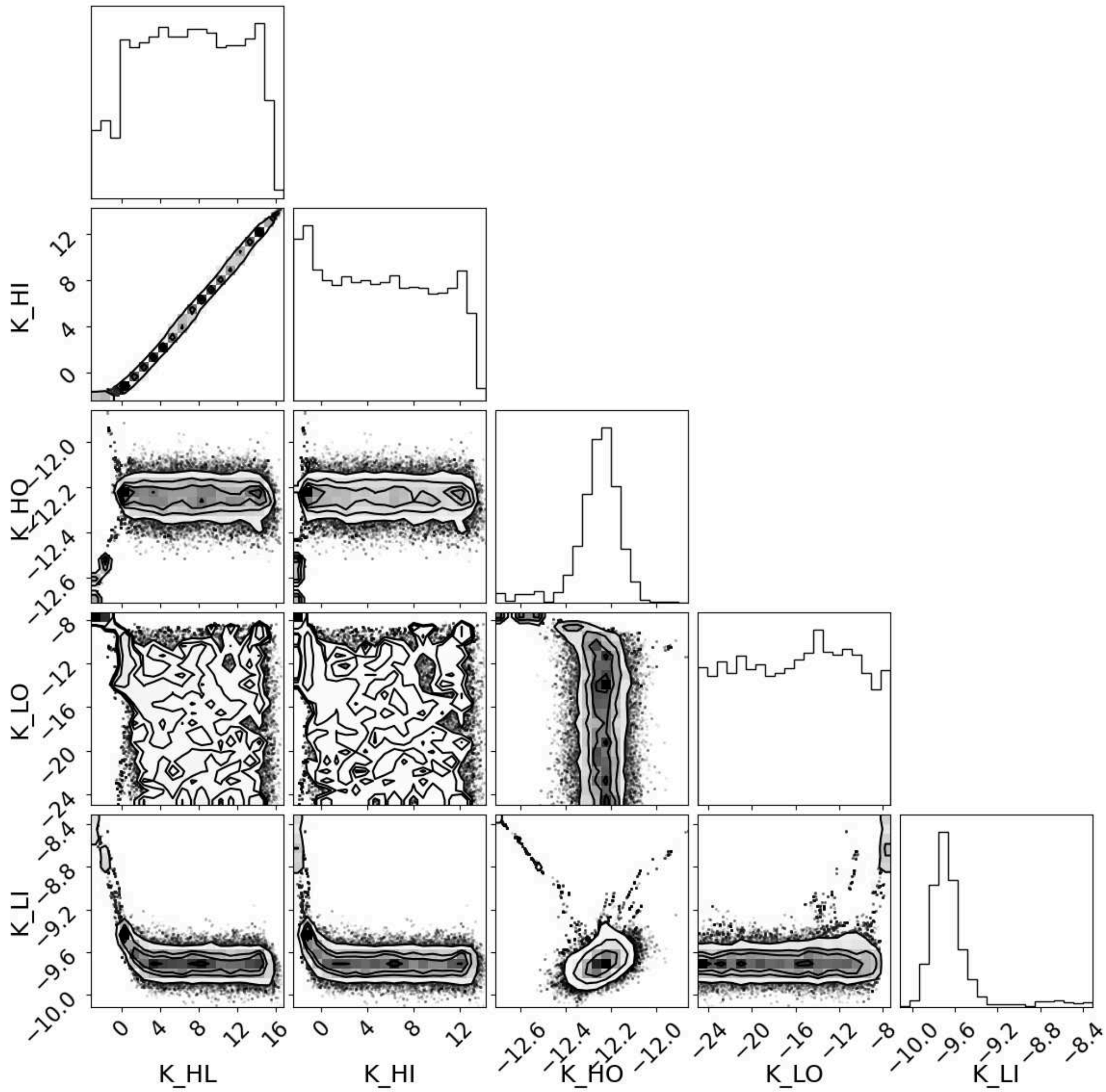

**Fig S10: Corner plot for MCMC samples of parameters for the LacMWC4 model against the binding data for the mhk genotype.** Parameters: ln($K_{H \cdot I}$), ln($K_{L \cdot I}$), ln($K_{H \cdot DNA}$), ln($K_{HL}$). Plots show marginal distribution of values for each model parameter (panels on diagonal) and correlation of fit parameters (panels off-diagonal) across MCMC samples. Blue lines indicate median values. This is the model used in the analyses in this manuscript.

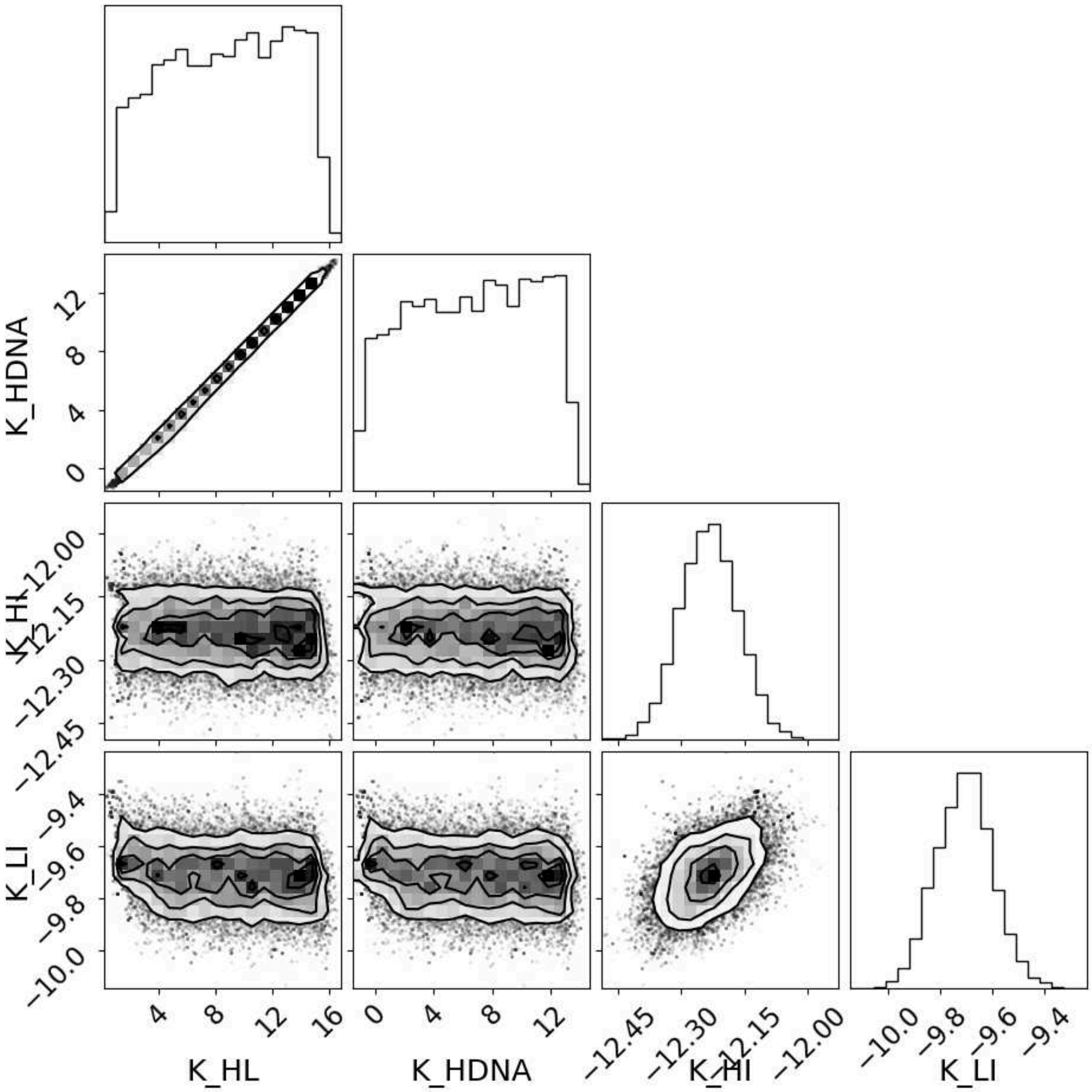

**Fig S11: Corner plot for MCMC samples of parameters for the LacMWC3i model against the binding data for the mhk genotype.** Parameters: $\ln(K_{H\cdot I})$, $\ln(K_{L\cdot I})$, $\ln(K_{H\cdot DNA})$, s. Plots show marginal distribution of values for each model parameter (panels on diagonal) and correlation of fit parameters (panels off-diagonal) across MCMC samples. Blue lines indicate median values. This is the model used in the analyses in this manuscript.

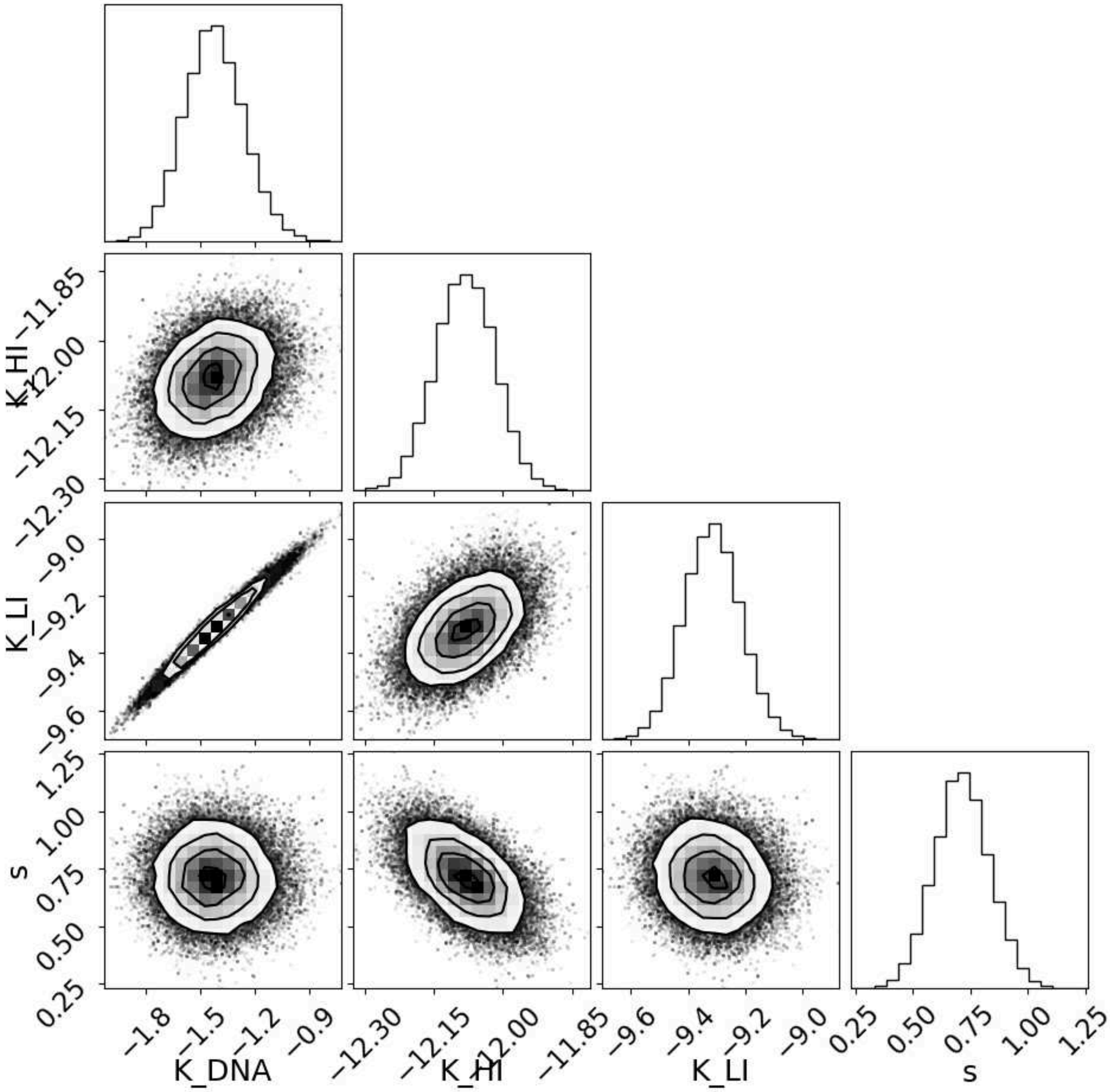

**Fig S12: Corner plot for MCMC samples of parameters for the LacMWC3 model against the binding data for the mhk genotype.** Parameters: $\ln(K_{H\cdot I})$, $\ln(K_{L\cdot I})$, $\ln(K_{H\cdot DNA})$. Plots show marginal distribution of values for each model parameter (panels on diagonal) and correlation of fit parameters (panels off-diagonal) across MCMC samples. Blue lines indicate median values.

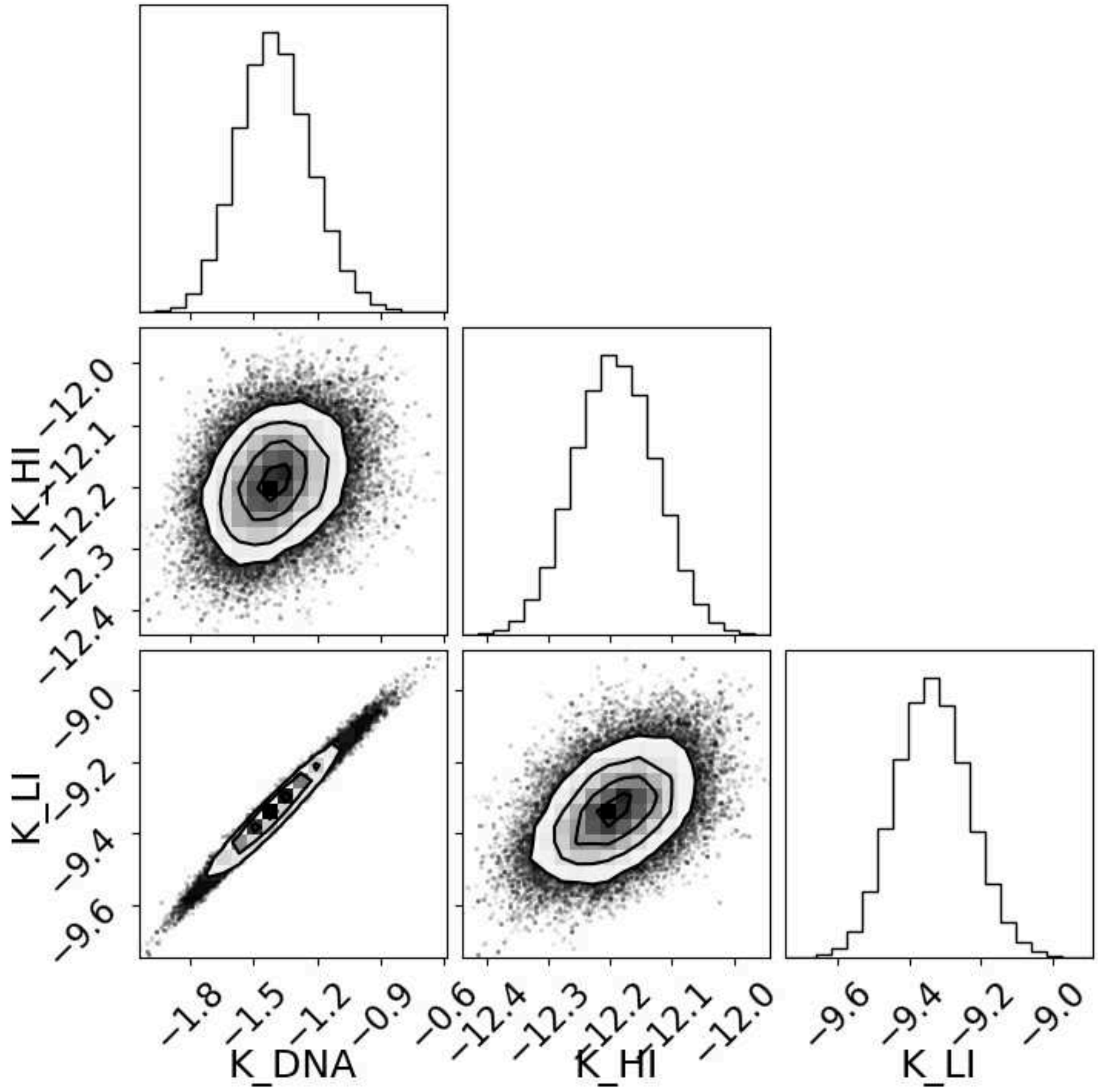

**Fig S13: Corner plot for MCMC samples of parameters for the LacMWC2 model against the binding data for the mhk genotype.** Parameters: $\ln(K_{H \cdot I})$ and $\ln(K_{H \cdot DNA})$. Plots show marginal distribution of values for each model parameter (panels on diagonal) and correlation of fit parameters (panels off-diagonal) across MCMC samples. Blue lines indicate median values.

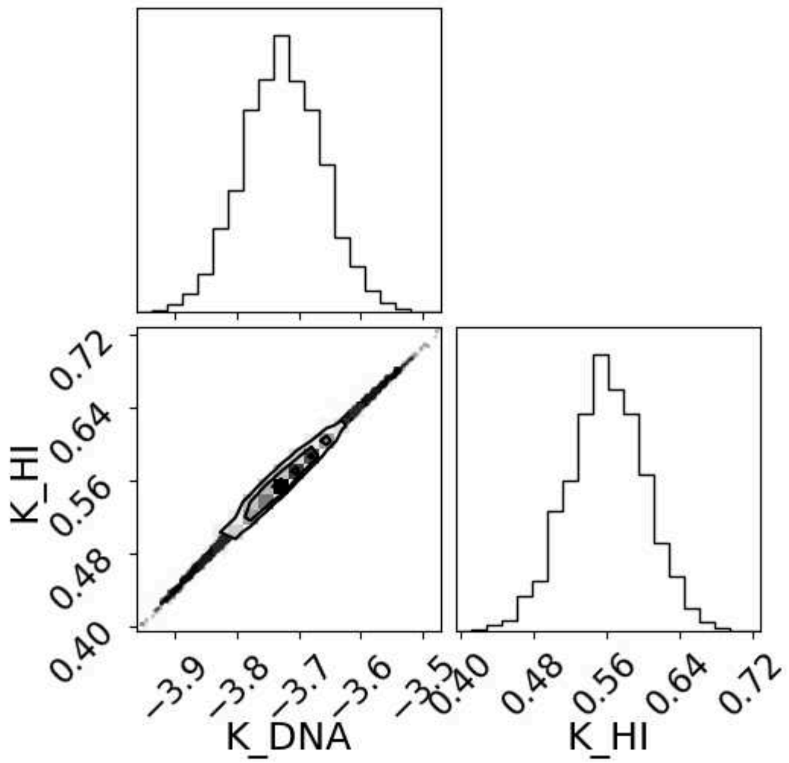

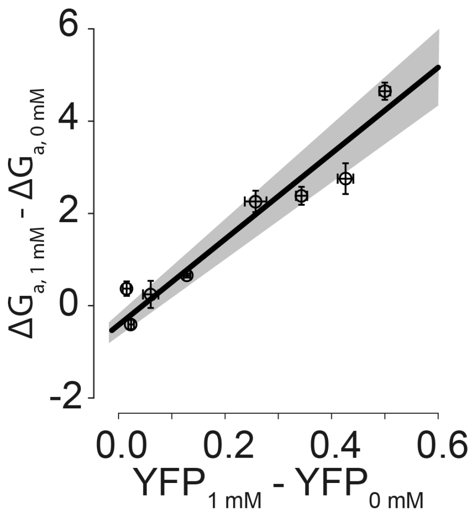


**Figure S14. IPTG-dependent change in ΔGa correlates with IPTG-dependent change in YFP expression.** Plot compares the change in binding energy between 0 and 1 mM IPTG ($\Delta\Delta G_a = \Delta G_{a,1mM} - \Delta G_{a,0mM}$) to the change in YFP induction between 0 and 1 mM IPTG ($YFP_{1mM} - YFP_{0mM}$). Each point is one of the eight genotypes studied. The errors reported on each $\Delta\Delta G_a$ value are propagated from the uncertainty from the MCMC samples; errors on YFP expression are propagated standard deviations from biological triplicates. The YFP induction and $\Delta\Delta G_{bind}$ values were strongly correlated ($R^2 = 0.88$).

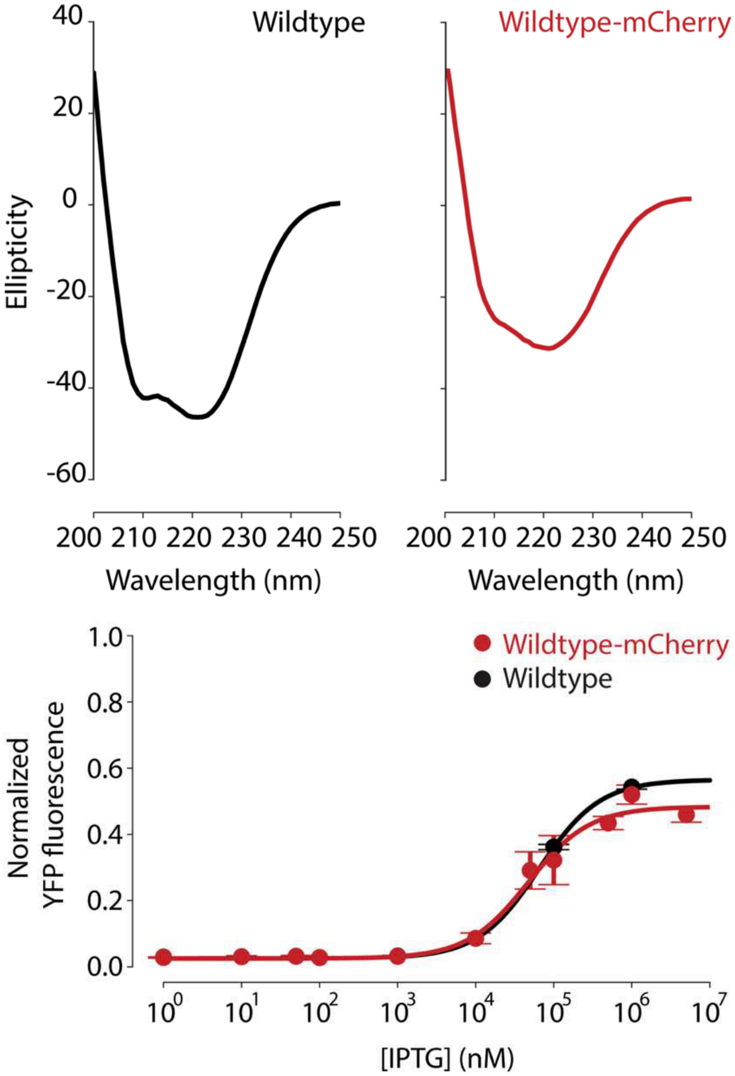


**Fig S15: mCherry-tagged lac repressor behaves similarly to the untagged repressor.** Top panel compares far-UV CD spectra of purified wildtype lac repressor and purified lac repressor fused to mCherry. Wavelength is indicated on the x-axis; raw ellipticity on the y-axis. Both proteins are folded. Bottom panel shows the YFP expression induced *in vivo* by the wildtype lac repressor (black) and the mCherry-tagged repressor (red) as a function of IPTG. YFP expression was measured as described in the text.

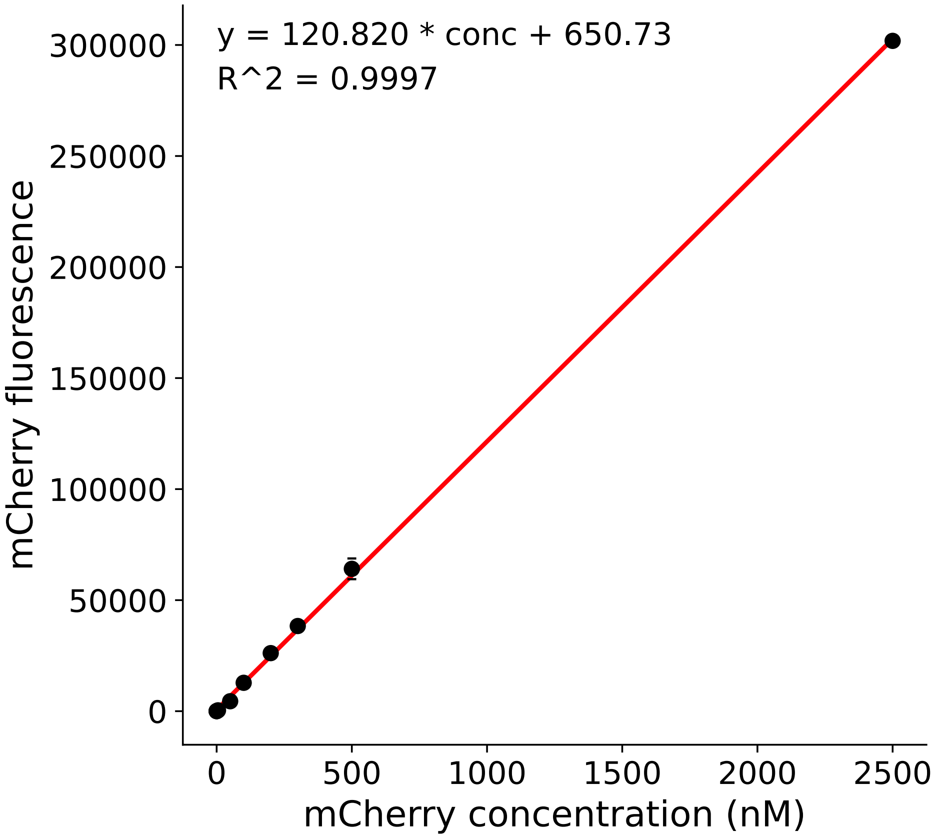


**Fig S16: Calibration curve relating mCherry concentration to mCherry fluorescence.** X-axis is mCherry protein concentration (serial dilution); y-axis is mean and standard deviation of mCherry fluorescence for each replicate for three biological replicates.

| variant | parameter | median | lower | upper | additive |
|---|---|---|---|---|---|
| mhk | $\Delta G^{\circ}_{DNA}$ | 0.92 | 0.70 | 1.12 | 0.92 |
| mhk | $\Delta G^{\circ}_{H\cdot I}$ | 7.76 | 7.68 | 7.84 | 7.76 |
| mhk | $\Delta G^{\circ}_{L\cdot I}$ | 5.98 | 5.85 | 6.11 | 5.98 |
| mhk | s | 2.04 | 1.62 | 2.59 | n/a |
| Ihk | $\Delta G^{\circ}_{DNA}$ | 0.39 | 0.10 | 0.62 | 0.39 |
| Ihk | $\Delta G^{\circ}_{H\cdot I}$ | 7.51 | 6.97 | 8.00 | 7.51 |
| Ihk | $\Delta G^{\circ}_{L\cdot I}$ | 6.56 | 6.06 | 7.01 | 6.56 |
| Ihk | s | 0.28 | 0.20 | 0.41 | n/a |
| mAk | $\Delta G^{\circ}_{DNA}$ | 2.10 | 1.74 | 2.41 | 2.10 |
| mAk | $\Delta G^{\circ}_{H\cdot I}$ | 4.92 | 3.00 | 6.60 | 4.92 |
| mAk | $\Delta G^{\circ}_{L\cdot I}$ | 3.82 | 1.89 | 5.50 | 3.82 |
| mAk | s | 2.35 | 1.19 | 4.46 | n/a |
| mhL | $\Delta G^{\circ}_{DNA}$ | 2.20 | 2.19 | 2.21 | 2.20 |
| mhL | $\Delta G^{\circ}_{H\cdot I}$ | 4.96 | 2.83 | 6.43 | 4.96 |
| mhL | $\Delta G^{\circ}_{L\cdot I}$ | 4.62 | 2.49 | 6.08 | 4.62 |
| mhL | s | 1.62 | 1.39 | 1.90 | n/a |
| IAk | $\Delta G^{\circ}_{DNA}$ | 0.48 | 0.17 | 0.74 | 1.57 |
| IAk | $\Delta G^{\circ}_{H\cdot I}$ | 5.56 | 4.61 | 6.25 | 4.67 |
| IAk | $\Delta G^{\circ}_{L\cdot I}$ | 4.60 | 3.63 | 5.31 | 4.39 |
| IAk | s | 0.36 | 0.27 | 0.47 | n/a |
| IhL | $\Delta G^{\circ}_{DNA}$ | 0.38 | 0.27 | 0.49 | 1.67 |
| IhL | $\Delta G^{\circ}_{H\cdot I}$ | 4.19 | 2.67 | 6.04 | 4.72 |
| IhL | $\Delta G^{\circ}_{L\cdot I}$ | 4.58 | 3.04 | 6.49 | 5.20 |
| IhL | s | 0.22 | 0.18 | 0.29 | n/a |
| mAL | $\Delta G^{\circ}_{DNA}$ | 2.62 | 2.28 | 2.96 | 3.39 |
| mAL | $\Delta G^{\circ}_{H\cdot I}$ | 4.74 | 1.96 | 8.06 | 2.13 |
| mAL | $\Delta G^{\circ}_{L\cdot I}$ | 4.59 | 1.81 | 8.02 | 2.46 |
| mAL | s | 2.90 | 2.02 | 4.46 | n/a |
| IAL | $\Delta G^{\circ}_{DNA}$ | 0.90 | 0.72 | 1.06 | 2.86 |
| IAL | $\Delta G^{\circ}_{H\cdot I}$ | 4.80 | 2.50 | 7.48 | 1.88 |
| IAL | $\Delta G^{\circ}_{L\cdot I}$ | 4.58 | 2.28 | 7.28 | 3.03 |
| IAL | s | 0.62 | 0.45 | 0.93 | n/a |

**Table S1: Parameter values for the lac repressor ensemble LacMWC3i model.** The “lower” and “upper” columns note the edges of the 95% credibility interval for the parameter across MCMC samples. The “additive” column indicates the parameter values used to generate the additive thermodynamic ensembles in Figure 5.

| model | $\ln(K_{HL})$ | $\ln(K_{H\cdot I})$ | $\ln(K_{L\cdot I})$ | $\ln(K_{H\cdot DNA})$ | $\ln(K_{L\cdot DNA})$ | s | $\ln\mathscr{L}$ | k | AIC weight |
|---|---|---|---|---|---|---|---|---|---|
| LacMWC2 | 0 | f | 0 | f | -23.0 | 1.0 | -8093 | 17 | $< 10^{-308}$ |
| LacMWC3 | 0 | f | f | f | -23.0 | 1.0 | 488.1 | 25 | $3 \times 10^{-63}$ |
| LacMWC3i | 0 | f | f | f | -23.0 | f | 633.1 | 33 | 1.000 |
| LacMWC4 | f | f | f | f | -23.0 | 1.0 | 347.6 | 33 | $1 \times 10^{-124}$ |
| LacMWC5 | f | f | f | f | f | 1.0 | -393.5 | 41 | $< 10^{-308}$ |

**Table S2: Ensemble model selection.** Table gives the floating parameters, log-likelihoods, and final AIC weight for each model. The model names match the function names from the software (https://github.com/harmslab/lacmwc). Model parameters set to "f" varied during MCMC sampling, while those with numerical values (e.g. $\ln(K_{HL}) = 0$) were fixed to that value. "$\ln\mathscr{L}$" gives the log likelihood of the median parameter estimates, summed over all datasets. "k" gives the number of fit parameters (number of floating parameters × eight variants + 1). The analyses reported in the manuscript used the LacMWC3i model.
